# Optical Imaging of Metabolic Dynamics in Animals

**DOI:** 10.1101/285908

**Authors:** Lingyan Shi, Chaogu Zheng, Yihui Shen, Zhixing Chen, Edilson S. Silveira, Luyuan Zhang, Mian Wei, Chang Liu, Carmen de Sena-Tomas, Kimara Targoff, Wei Min

## Abstract

Direct visualization of metabolic dynamics in living tissues with high spatial and temporal resolution is essential to understanding many biological processes. Here we introduce a platform that combines <u>d</u>euterium <u>o</u>xide (D_2_O) probing with <u>s</u>timulated <u>R</u>aman <u>s</u>cattering microscopy (DO-SRS) to image *in situ* metabolic activities. Enzymatic incorporation of D_2_O-derived deuterium into macromolecules generates carbon-deuterium (C-D) bonds, which track biosynthesis in tissues and can be imaged by SRS *in situ*. Within the broad vibrational spectra of C-D bonds, we discovered lipid-, protein-, and DNA-specific Raman shifts and developed spectral unmixing methods to obtain C-D signals with macromolecular selectivity. DO-SRS enabled us to probe *de novo* lipogenesis in animals, image protein biosynthesis without tissue bias, and simultaneously visualize lipid and protein metabolism and reveal their different dynamics. DO-SRS, being noninvasive, universally applicable, and cost-effective, can be adapted to a broad range of biological systems to study development, tissue homeostasis, aging, and tumor heterogeneity.

## Introduction

Understanding the dynamics of metabolism in multicellular organisms is important to unraveling the mechanistic basis of many biological processes, such as development, aging, energy homeostasis, and tumor progression and heterogeneity. Although metabolomic technologies can catalog thousands of metabolites residing in cells, nondestructive tools are limited for *in situ* visualization of metabolic activities, such as protein and lipid synthesis and degradation, at subcellular resolution in living organisms. Magnetic resonance spectroscopic imaging (MRSI) and positron emission tomography (PET) can provide metabolic information noninvasively and have wide oncological application but lack sufficient spatial resolution^1^.Microautoradiography and fluorescence microscopy can visualize metabolism at the single-cell level but require radioactive and fluorescent labeling of the substrate, respectively; these labeling are often toxic to cells and often perturb the native metabolic processes^2^. Nanoscale Secondary Ion Mass Spectrometry (NanoSIMS) and the more recent multi-isotope imaging mass spectrometry (MIMS) can measure the incorporation of nontoxic stable isotopes, like ^15^N and ^13^C, at submicrometer resolution and spatially track the labeling of biomolecules; but both methods are destructive to living tissues and have limited resolvability for macromolecules^3, 4^.

Here we developed a general method that combines <u>d</u>euterium <u>o</u>xide probing and <u>s</u>timulated <u>R</u>aman <u>s</u>cattering microscopy (DO-SRS) to provide imaging contrast for visualizing metabolic dynamics *in situ*. Through systematic investigation of the carbon-deuterium (C-D) vibrational spectrum, we discovered Raman shifts associated with C-D bond-containing lipids, proteins, and DNA, respectively, and further revealed that this spectral selectivity resulted from the sparse labeling pattern and inherently different chemical environments surrounding the C-D bond in different types of macromolecules. By applying DO-SRS to living cells and animals, we not only demonstrated its broad utility, high sensitivity, noninvasiveness, subcellular resolution, compatibility with other imaging modality, and suitability for *in vivo* live imaging in mammals, but also gained new insights on the metabolic basis of several biological processes.

## Results

### SRS imaging enables D_2_O to be a contrast agent for metabolic activities

Water (H_2_O), the ubiquitous solvent of life, diffuses freely across cell and organelle membranes and participates in the vast majority of biochemical reactions. As an isotopologue of water, heavy water (D_2_O) can rapidly equilibrate with total body water in all cells within an organism and label cellular biomolecules with deuterium (D) by forming a variety of X-D bonds through non-enzymatic H/D exchange and enzymatic incorporation (Figure 1A). The former is spontaneous and reversibly forms oxygen-deuterium (O-D), nitrogen-deuterium (N-D), and sulfur-deuterium (S-D) bonds on existing molecules, whereas the latter depends on enzyme-catalyzed chemical transformation that irreversibly breaks the O-D bond and forms C-D bonds on newly synthesized molecules^5^. Through such transformation, deuterium quickly labels the metabolic precursors, such as non-essential amino acids (NEAAs), acetyl-CoA, and deoxyribose, which are then slowly incorporated into proteins, lipids, and DNA, respectively^6-8^ (Figure 1A). As often the rate-limiting step, the synthesis of C-D bond-containing macromolecules from the precursors is governed by cellular metabolic activities. Therefore, D_2_O can be used as a universal probe to track metabolic rate through the emergence of C-D bond-containing macromolecules (hereinafter referred to as D-labeled macromolecules).

**Figure 1.**
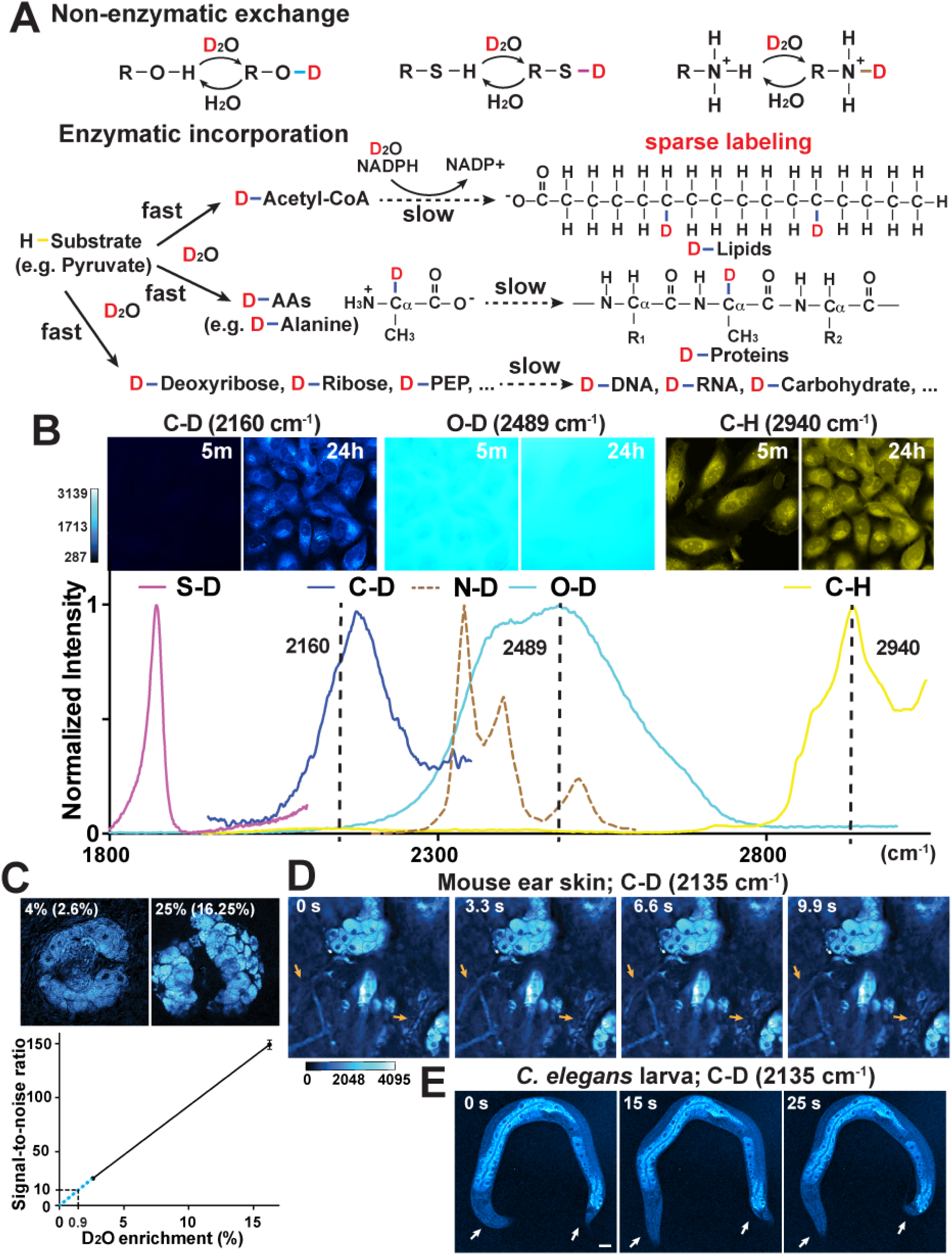
SRS imaging of biosynthetic incorporation of deuterium into macromolecules in living cells and animals. (A) D_2_O-derived deuterium can form O-D, S-D, and N-D bonds through reversible non-enzymatic H/D exchange and be incorporated into C-D bonds of metabolic precursors for the synthesis of macromolecules through irreversible enzymatic incorporation. (B) Various X-D bonds produced Raman peaks at distinct positions. C-D and C-H spectra were collected from Hela cells grown in DMEM containing 70% D_2_O for 24 hours, O-D spectrum from the 70% D_2_O medium, S-D spectrum from saturated cysteine dissolved in D_2_O, and N-D spectrum adopted from^60^. In the top row, SRS images were collected for C-D, O-D, and C-H signal of HeLa cells at 5 minutes and 24 hours after adding D_2_O-containing medium. (C) Signal-to-noise ratio (S/N; noise = 1 μV) for SRS signal of sebaceous glands at 2135 cm^-1^ from mice that drank 4% or 25% D_2_O for 8 days. Percentages of D_2_O enrichment in body water are shown in parentheses. Detection limit at S/N = 10 or 2 were calculated based on the linear relationship between average S/N and D_2_O enrichment level. (D-E) Frames from live SRS imaging recordings of the sebaceous glands under the ear skin of intact mice that drank 25% D_2_O for 9 days (Movie S1-S2) and living fourth stage *C. elegans* larva that grew on 20% D_2_O-containing NGM plates for 4 hours (Movie S3-S4). The blood flow (orange arrows) in mouse visualized with two-photon absorption contrast and the movement of the *C. elegans* body (white arrows) indicated that the animals under imaging were alive.

Raman spectroscopy provides a noninvasive, optical approach to distinguish metabolic incorporation from non-enzymatic exchange *in situ*, because various X-D bonds have intrinsically distinct stretching vibrational features. We found that the Raman spectrum of C-D bond was clearly separated from those of C-H, O-D in D_2_O, and the non-enzymatically formed O-D, S-D and N-D bonds (Figure 1B), which allows the direct detection of biosynthetic incorporation of deuterium through the amount of C-D bonds. In fact, spontaneous Raman microspectroscopy has recently been employed to identify metabolic activity in bacteria after D_2_O treatment^9, 10^. However, although gaining popularity in microbiology, this approach has difficulties in generating spatially resolved images due to low sensitivity and slow imaging speed. Furthermore, it only measures the total intensity of C-D bonds without distinguishing the types of macromolecules. In practice, this method has only been applied to single-cell organisms with high metabolic activity, but not metazoans that may have slower metabolism, hence weaker signal, and require higher detection sensitivity.

Compared to spontaneous Raman spectroscopy, stimulated Raman scattering (SRS) microscopy is an emerging nonlinear Raman imaging technology with substantial sensitivity boost through quantum amplification by stimulated emission, which enables at least three orders of magnitude faster acquisition time, fine spectral resolution, compatibility with fluorescence, and 3D optical sectioning capability in tissues and even living animals^11, 12^. These unique advantages of SRS microscopy have rendered it the most powerful vibrational imaging technique with expanding impact to biophotonics^13-16^. Combined with our new discoveries of the chemical features of the C-D vibrational spectrum (described below), DO-SRS will allow the development of D_2_O into a powerful imaging contrast agent that can specifically trace lipid, protein, and DNA metabolism in cells and tissues.

We first demonstrated DO-SRS imaging on the metabolism of cultured cells. By treating HeLa cells with medium containing 70% D_2_O for 24 hours (we quantified ATP production by metabolically active cells and found that toxicity only arises when D_2_O concentration exceeded 80%; Supplementary Figure 1) and then tuning SRS to target the C-D frequency, we found that C-D signal was undetectable at the beginning of the treatment but increased dramatically in all cells at 24 hours (Figure 1B). This result confirms that C-D signal specifically and effectively reports newly synthesized molecules, whereas C-H signal represents pre-existing pool of molecules. Importantly, the separation between the O-D peak and the C-D peak means that C-D signal is essentially free of interference from the overwhelming O-D background, thus washing off the D_2_O probe before imaging is unnecessary.

### Safety, sensitivity, and *in vivo* live imaging capacity of DO-SRS

In humans, deuterium as a stable isotope is widely used to measure body composition and metabolic rate^17-19^. Daily intake of 60 to 70 ml D_2_O, which results in ∼2% body water enrichment, does not cause any adverse symptoms^20, 21^, and is considered to be safe. We found that a comparable level (2.4∼2.8%) of enrichment in mice by administration of 4% D_2_O as drinking water (the dilution of body water relative to drinking water is typically 30∼40% in rodents; ^21^) produced easily detectable C-D signals in sebaceous glands (Figure 1C).

In mice, D_2_O enrichment below 20% did not cause any effect on physiological processes, including no acute adverse events, no effects on cell division in all the major cell renewal systems, no perturbation on physiology, growth, appetite, and reproduction, and no teratogenic effects, even in multigenerational studies^19, 22, 23^. Thus, we gave mice 25% D_2_O as drinking water to achieve a safe level (15∼17.5%) of enrichment in body water and obtained a 6-fold increase in C-D signal compared to that from mice drinking 4% D_2_O (Figure 1C). The nearly linear relationship between D_2_O enrichment and signal intensity allowed us to extrapolate the detection limit (Figure 1C). Administration of 1.4% D_2_O (∼0.9% enrichment) is sufficient to achieve a signal-to-noise ratio (S/N) of 10, and for S/N higher than 2, administration of 0.3% D_2_O would be needed. Although detectable signal can be generated at low D_2_O enrichment, we chose ∼16% D_2_O enrichment (from oral administration of 25% D_2_O) in the following mouse studies in order to achieve large dynamic range and also reveal low biosynthetic activity within a short time frame.

The wash-free aspect of DO-SRS enables us to image the real-time C-D signal dynamics of live cells and intact animals in the presence of D_2_O. We performed *in vivo* live imaging of the sebaceous glands under the ear skin of anesthetized mice that drank 25% D_2_O for 9 days. Leveraging the 3D optical sectioning and substantial imaging depth of nonlinear excitation of SRS (about hundreds of microns), we observed clear C-D signal in sebocytes in living mice (Figure 1D and Movie S1-3). To our knowledge, this is the first time that deuterium-labeled macromolecules have been visualized in intact living mammals, thanks to the high-speed live imaging capability of SRS (> 1000 times faster than spontaneous Raman microspectroscopy). At a whole-organism level, we were able to monitor C-D signals in moving *C. elegans* larva growing in a 20% D_2_O environment (Figure 1E and Movie S4-5). In *C. elegans*, < 60% D_2_O concentration had no observable toxicity and did not affect cell division (Supplementary Figure 1). With the above proof of concept, we show that the safeness of D_2_O as a probe, combined with the sensitivity and penetration of SRS as a detection mechanism, would allow for long-term *in vivo* tracking of metabolic activities.

### Identification of macromolecule-specific frequencies in C-D vibrational spectra

Because D_2_O is a universal probe and enables the labeling of different types of macromolecules, the C-D signal generated from D_2_O probing is considered to report the total metabolic activity. Indeed, earlier Raman spectroscopy studies simply used the peak intensity to represent the amount of deuterium incorporation into total biomass^9^. Whether sufficiently distinct C-D vibrational frequencies exist to allow the separation of different types of D-labeled macromolecules is unaddressed.

To address this question, one might attempt to translate the spectral knowledge of C-H region to the C-D region, but we reasoned that this translation does not apply. C-H stretching vibrational spectra contain a main peak at 2940 cm^-1^ (denoted as CH_*P*_ channel) originating from protein-related CH_3_ stretching, a main peak at 2845 cm^-1^ (CH_*L*_ channel) originating from lipid-related CH_2_ stretching (Fig 1B), and a shoulder peak at 2967 cm^-1^ (CH_*DNA*_ channel) from DNA-related C-H stretching; those three Raman shifts were used to map endogenous cellular lipids, proteins, and DNA in label-free SRS^24-26^. However, the C-H spectral distinction (based on the number of covalent hydrogen atoms) cannot be simply translated to C-D stretching, because CD_2_ and CD_3_ groups would rarely exist under relatively low D_2_O enrichment (15∼17.5%) in body water. Such low D_2_O concentration means that deuterium could only sparsely label the newly synthesized macromolecules (Figure 1A), and the C-D vibrational signal would be dominated by CD mode (only one carbon-bonded H atom is replaced by D atom).

We set out to dissect the spectral features of C-D vibrational modes upon D_2_O treatment and then to investigate the underlying chemical basis. To begin with, we scanned the C-D region (2109-2210 cm^-1^), tested 5 Raman shifts with equal intervals, and found identifiable difference among the 5 images (Figure 2A). C-D images acquired at lower wavenumbers (especially 2135 cm^-1^) resembled the cellular distribution patterns of lipids from the CH_*L*_ channel, and images acquired at higher wavenumbers (especially 2185 cm^-1^) resembled the protein signal from CH_*P*_ channel in all tissues we tested, including fibroblast-like COS7 cells, *C. elegans* larva, zebrafish embryos, and mouse tissues (Figure 2A). Thus, we hypothesized the signal at 2135 cm^-1^ may represent newly synthesized, D-labeled lipids and the signal at 2185 cm^-1^ as D-labeled proteins.

**Figure 2.**
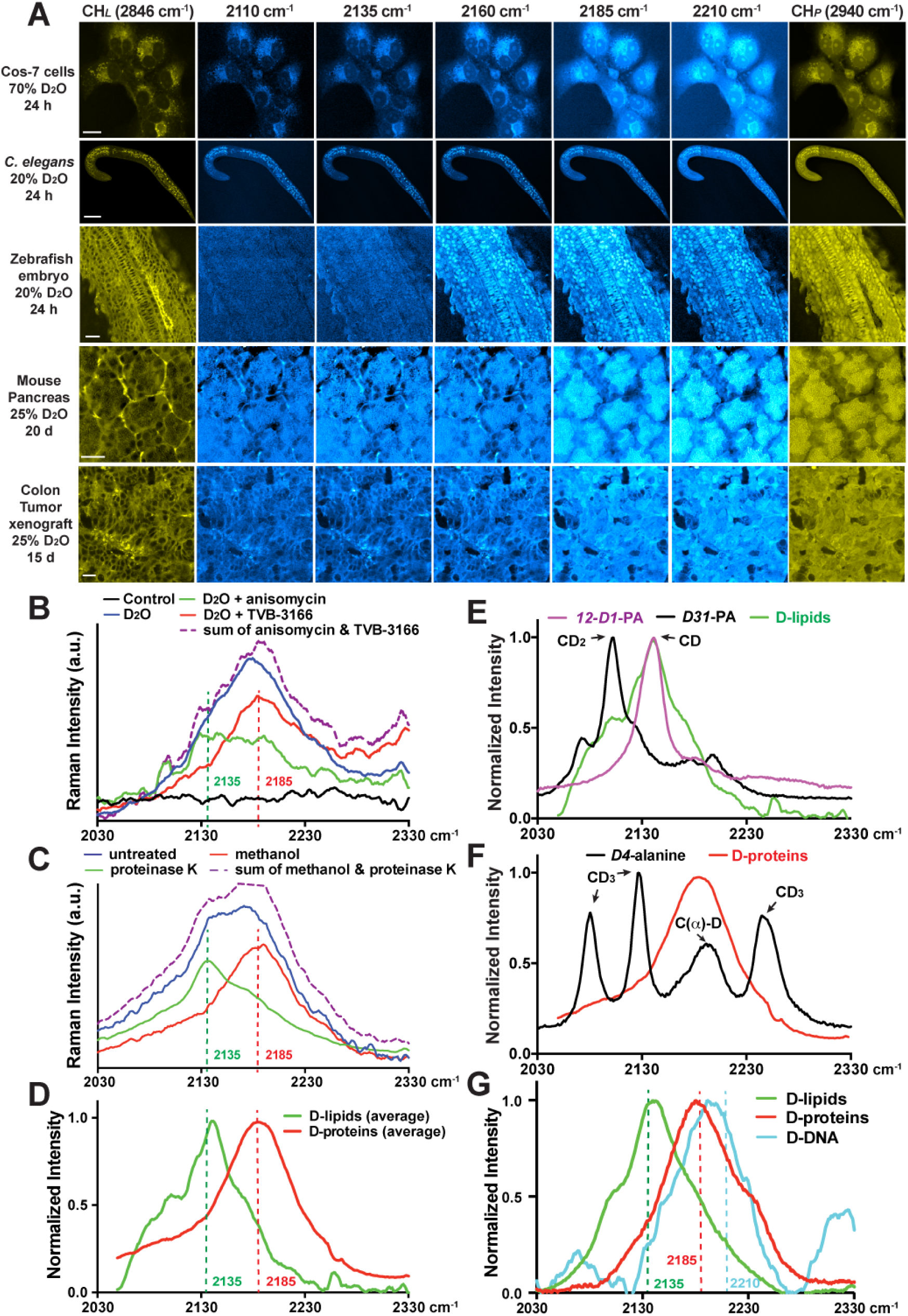
Identification of specific Raman shifts with macromolecular selectivity within the broad C-D vibrational spectra. (A) SRS images of various cells and tissues from animals treated with D_2_O for indicated amounts of time. Images were collected using previously known Raman shifts for CH-containing lipids (CH_*L*_ 2846 cm^-1^) and proteins (CH_*P*_ 2940 cm^-1^) and five wavenumbers (2110, 2135, 2160, 2185, and 2210 cm^-1^) within the C-D broadband. Scale bar = 20 μm. (B) Spontaneous Raman spectrum of Hela cells grown in DMEM made of 70% D_2_O in the absence or presence of fatty acid synthase inhibitor TVB-3166 or protein synthesis inhibitor anisomycin. Cells grown in DMEM made of 100% H_2_O were used as control (black). Purple curve shows the sum of the values on the green and red curves. (C) Spontaneous Raman spectrum of deuterium-labeled xenograft colon tumor tissues treated with protease K or washed with methanol for 24 hours. Mice bearing the xenograft drank 25% D_2_O as drinking water for 15 days before tumor tissues were harvested and imaged. (D) The normalized Raman spectra of tissue after 24-hour methanol wash (D-labeled protein signal in red) and the difference spectra before and after methanol wash (D-labeled lipid signal in green), averaged from various mouse tissues. (E) Comparison of Raman spectra of *12-D1*-palmitic acid (100 mM dissolved in DMSO), *D31*-palmitic acid (100 mM in DMSO), and *in situ* D-labeled lipid standards. (F) Comparison of Raman spectra of *D4*-alanine (100 mM in PBS), and D-labeled protein standards. Assignment of the peaks were made according to a previous report^27^. (G) Raman spectra of biochemically extracted lipids, proteins, and DNA from HeLa cells grown in DMEM media containing 70% D_2_O.

To confirm this hypothesis, we first added anisomycin to inhibit protein synthesis while growing HeLa cells in D_2_O-containing medium and found the Raman spectrum had a peak centered at 2135 cm^-1^, representing the remaining D-labeled lipid signal (Figure 2B). Conversely, blocking lipid synthesis with fatty acid synthase inhibitor TVB-3166 led to a peak centered at 2185 cm^-1^, representing the remaining D-labeled protein signal (Figure 2B). The sum of the signals from the two treatments appeared similar to the control group treated only with D_2_O. At the tissue level, removing proteins by proteinase K treatment in mouse tissues abolished the peak at 2185 cm^-1^, and dissolving lipids by methanol abolished the peak at 2135 cm^-1^ (Figure 2C). Applying methanol wash to multiple mouse tissues from various organs, we obtained the average spectra for D-labeled lipids (signal reduction by methanol wash) and proteins (residual signal after methanol wash), consistently showing peaks at the two frequencies (Figure 2D). The above data confirmed macromolecule-specific Raman shifts for D-labeled lipids and proteins at 2135 cm^-1^ (denoted as CD_*L*_ channel) and 2185 cm^-1^ (CD_*P*_ channel), respectively.

We then sought for the chemical basis of spectral distinction in CD vibration by comparing D-labeled lipid and protein spectra with assigned CD vibrational modes in model compounds. We found that the peak of D-labeled lipid at ∼2140 cm^-1^ matched well with the singly deuterated C-D stretching in *12-D1*-palmitic acid but not the CD_2_ symmetric stretching mode at ∼2100 cm^-1^ in perdeuterated palmitic acid (*D31*-) (Figure 2E), supporting the idea of sparse labeling (Figure 1A). The broadening of the peak in D-labeled lipids compared to *12*-*D1*-palmitic acid reflected a much more heterogeneous environment for the C-D bonds in lipids *in vivo*. Similarly, comparing to the spectrum of *D4*-alanine, we found that the D-labeled protein peak around 2185 cm^-1^ matched the peak for C(*α*)-D vibration but not the other three peaks assigned to the side chain CD_3_ group^27^ (Figure 2F). This pattern also supported sparse labeling, and moreover, indicated that most of the deuterium labeling in newly synthesized proteins occurred at the *α* carbon position through reversible transamination of free AAs^28^ (Figure 1A).

The above evidence suggests that the underlying principle for the observed spectral separation of D-labeled macromolecules is their distinct chemical microenvironments. Specifically, we attribute the spectral separation between D-labeled lipids (around 2140 cm^-1^) and proteins (around 2185 cm^-1^) to the inherently different chemical environments of the constituting fatty acids and amino acids—deuterium-bonded carbon atoms in a hydrocarbon chain of lipids and the main chain of polypeptides are connected to chemical groups with different polarities. We next asked if such principle can be extended to other macromolecules. Thus, we chemically isolated the major macromolecules from D_2_O-treated HeLa cells, including lipids, proteins, and DNA. As expected, the Raman spectra of D-labeled lipids and proteins matched well with the corresponding spectra obtained *in situ*. Importantly, we also obtained the spectrum of extracted DNA and found that the Raman peak of D-labeled DNA was blue-shifted compared to D-labeled proteins (Figure 2G). Deuterium labels DNA with higher chance at the C1’ and C2’ positions on the deoxyribose ^21, 29^, and the blue shift of DNA’s CD peak may be attributed to the fact that more carbon atoms in deoxyribose are bonded to electronegative oxygen atoms. Based on the spectra of the cellular extracts, we chose to image DNA at 2210 cm^-1^ (designated as CD_*DNA*_ channel).

In summary, we uncovered specific Raman shifts that identify particular types of D-labeled macromolecules, as well as sparse labeling and chemical microenvironment as the underlying chemical basis. Upon D_2_O treatment, this macromolecular selectivity would enable the separation of CD signals into distinct channels that reflect the dynamics of different metabolic pathways.

### Spectral unmixing of D-labeled lipid, protein, and DNA signals

Although the peaks of CD signals in lipids, proteins, DNA are separated, their overall spectra overlap substantially, suggesting that in a given channel for one type of macromolecules there may be bleed-through signals from the other two types (Figure 2D and G). Thus, we next developed a three-component unmixing algorithm to computationally decompose the mixed CD signals into three macromolecule-specific elements, removing the bleed-through signal for each channel and revealing the accurate spatial distribution for each type of D-labeled macromolecules.

We used chemically extracted, D-labeled total lipids, proteins, and DNA as pure standards to calculate a set of unmixing coefficients (Supplementary Figure 2A and Methods). Applying the three-component unmixing algorithm to dividing cells, we successfully separated the three types of D-labeled macromolecules. In particular, unmixing removed lipid and protein bleed-through in the CD_*DNA*_ channel and revealed clean D-labeled DNA signal that was only localized in the condensed chromosomes during mitosis (Figure 3A). Signal for D-labeled DNA was very weak in non-dividing cells due to the lack of DNA synthesis and the lower DNA density in the nucleus compared to mitotic cells. Given this fact and that *in vivo* DNA labeling can be achieved by other methods like BrdU staining, we did not focus on imaging the dynamics of DNA metabolism in this study. Instead, we focused on generating signals for D-labeled lipids and proteins, which are much more difficult to optically image *in vivo* using other methods.

**Figure 3.**
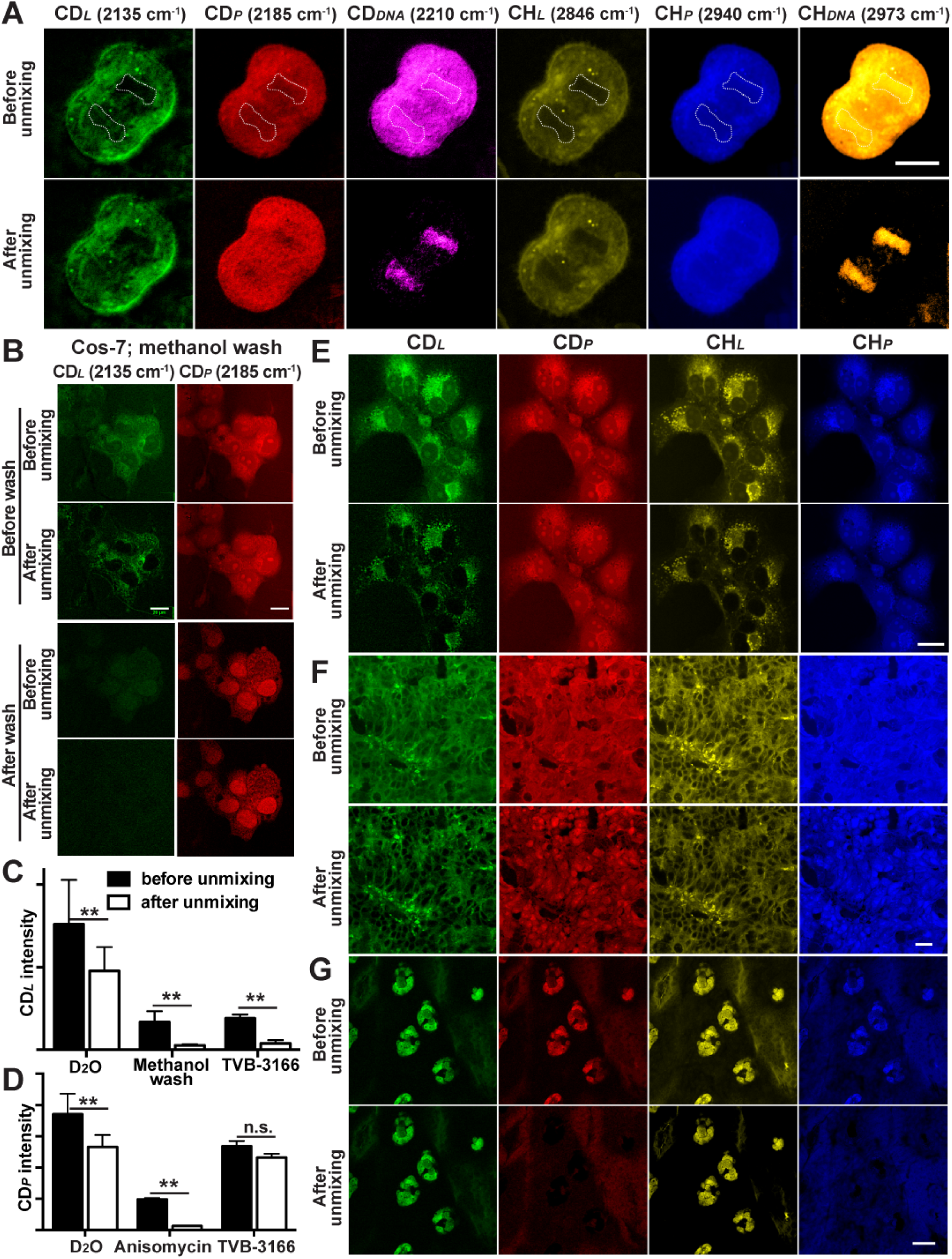
Spectral unmixing of D-labeled lipids, proteins, and DNA. (A) Separation of CD protein and DNA signals *via* unmixing in dividing cells (see Methods for details). Dashed outlines enclose the nuclei. In all Figures, CD_*L*_, CD_*P*_, CD_*DNA*_, CH_*L*_, CH_*P*_, and CH_*DNA*_ channels show signals collected at 2135, 2185, 2210, 2846, 2940, and 2973 cm^-1^, respectively, and are color-coded in green, red, pink, yellow, blue, and gold respectively. (B) SRS images, collected from the CD_*L*_ and CD_*P*_ channel, of COS-7 cells grown in 70% D_2_O DMEM for 24 hours and images of the same cells after methanol wash, with or without the application of unmixing algorithm. (C-D) Quantification of the mean SRS intensity (mean ± SD) from CD_*L*_ and CD_*P*_ channels of Cos-7 cells under various conditions before and after unmixing (N > 5 for each condition). ** indicates *p* < 0.01 in an unpaired t-test (also applied to other Figures). (E-G) Example sets of images before and after the application of CD_*L*_/CD_*P*_ unmixing for COS-7 cells grown in 70% D_2_O-containing DMEM for 24 hours (E), xenograft colon tumor tissues from mice drinking 25% D_2_O for 15 days (F), and sebaceous gland tissues from mice drinking 25% D_2_O for 3 days (G). CH_*L*_/CH_*P*_ unmixing was performed according to previous studies^24^. Scale bar = 20 μm. Tissue-specific unmixing parameters can be found in Methods.

To unmix CD_*L*_ and CD_*P*_ signals, we simplified the three-component unmixing algorithm to a two-component equation and applied it to images acquired at CD_*L*_ and CD_*P*_ channels. We validated the effectiveness of unmixing by showing that it abolished the residual bleed-through protein signal in the CD_*L*_ images of cells treated with TVB-3155 and the residual bleed-through lipid signal in the CD_*P*_ images of cells treated with anisomycin (Figure 3B-D). At the tissue level, we generated more accurate unmixing coefficients using the spectra of pure *in situ* D-labeled lipid and protein (lipid-free) signals obtained by methanol wash (Supplementary Figure 2B and C). We noticed some variation in the level of protein bleed-through into CD_*L*_ channel across different tissue types (Supplementary Figure 2C) and adjusted the coefficients accordingly (see Methods for details). Applying the proper calculation to tissue images, we completely removed the protein bleed-through (lipid signal after methanol wash) in the CD_*L*_ channel and revealed the genuine distribution of D-labeled lipids and proteins (Supplementary Figure 2D-F). For example, the nucleus became devoid of CD_*L*_ signal after unmixing in both cultured cells and tissues (Figure 3E and F). In lipid-rich tissues, unmixing effectively removed the strong lipid-to-CD_*P*_ bleed-through and revealed the true D-labeled protein signal (Figure 3G). Overall, our unmixing technique enables, for the first time, *in situ* deconvolution of D-labeled lipids and proteins signal *via* SRS microscopy.

It is worth noting the difference between the unmixing method developed above and the label-free counterpart previously reported. Unlike the unmixing of CH_*L*_ and CH_*P*_, which used pure standard compounds (oleic acid for lipids and Bovine Serum Albumin for proteins) ^24, 25^, our unmixing method is tailored for sparse labeling pattern of deuterium incorporation *in vivo*. Without commercially available standards for randomly, sparsely deuterated lipids and proteins, we generated standards using either chemically isolated or *in situ* D-labeled lipids and proteins to determine the unmixing coefficients.

Detailed understanding of the C-D spectral features and the development of unmixing methods allowed us to use DO-SRS to selectively probe *de novo* lipid and protein biosynthesis *in vivo*. The ability to separate the signals for D-labeled lipids and proteins enabled simultaneous visualization of the metabolic dynamics of lipid and protein in the same tissue, which is important for addressing many fundamental questions about the different pathways of cellular metabolism.

### Optical imaging of *de novo* lipogenesis *via* DO-SRS

Previously, our lab and others have developed methods to visualize lipogenic activities in living tissues by supplying deuterium-labeled fatty acids (D-FAs), such as palmitic acid, oleic acid, and arachidonic acid, and imaging C-D bonds in newly synthesized lipids^30-33^. However, DO-SRS is fundamentally different from those previous methods because D_2_O is a noninterfering probe that does not change native metabolism and is a non-carbon tracer that can probe activities of *de novo* lipogenesis.

D-FAs are known to be taken up by cells through scavenger pathways and then incorporated directly into lipids, whereas D_2_O freely diffuses into cells and labels newly synthesized lipids through *de novo* lipogenesis. Moreover, the dependence on cellular uptake for D-FA can also result in bias among various cell types, which does not occur for D_2_O probing. The different effects of these two types of probes are clearly evident in cultured cells. For example, supplementation of D-labeled palmitic acids (D-PA) or oleic acids (D-OA) in HeLa cells led to the accumulation of CD_*L*_ signal in lipid droplets (Figure 4A), whereas D_2_O probing generated very few lipid droplets in both CD_*L*_ and CH_*L*_ channels and produced much more uniform CD_*L*_ signal in cytoplasmic membrane structure.

**Figure 4.**
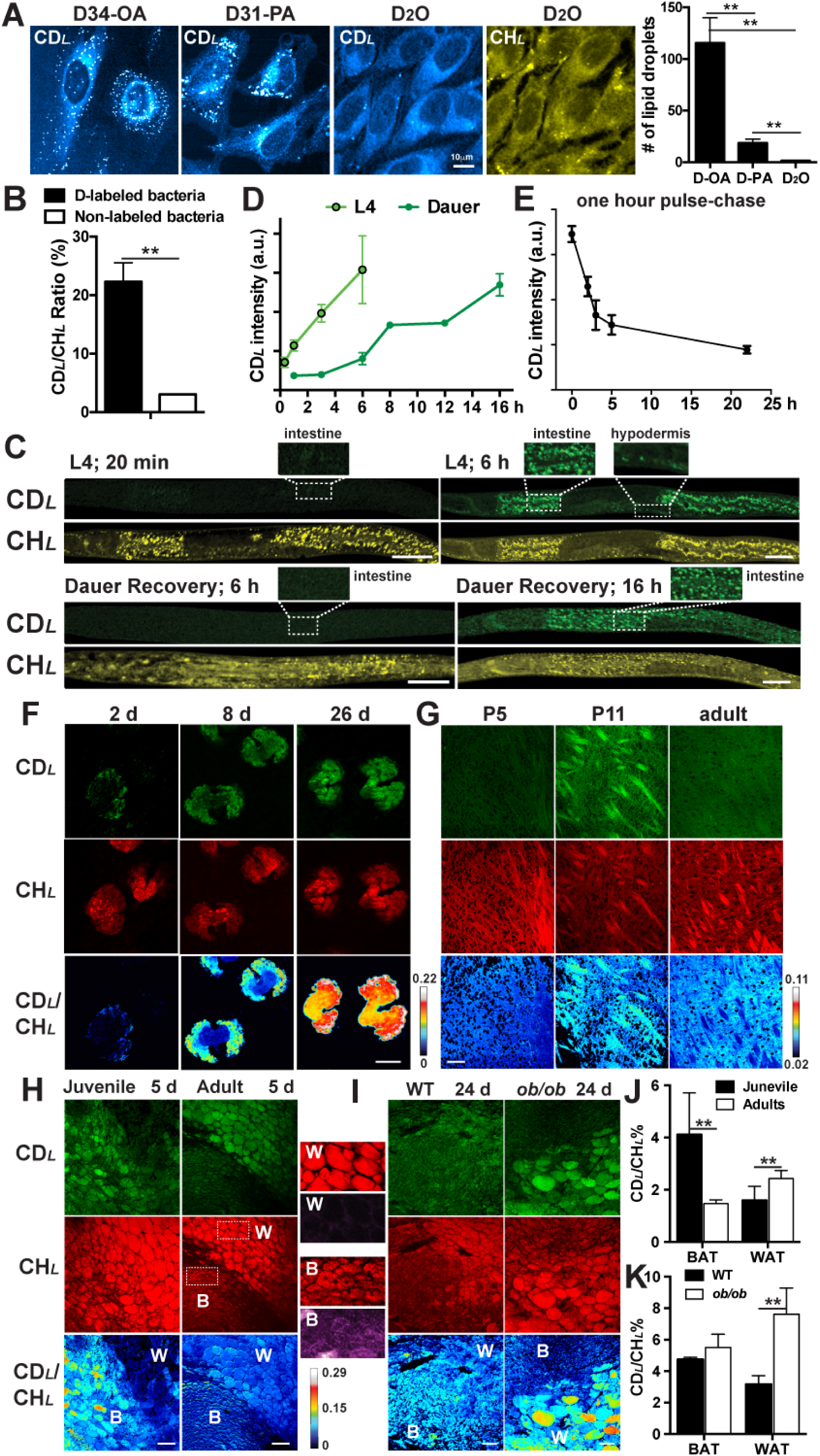
DO-SRS visualizes *de novo* lipogenesis *in vivo*. (A) SRS signals at 2110 cm^-1^ of HeLa cells grown in DMEM with 1% FBS and 10 μM of *D34*-oleic acids (D-OA) or *D31*-palmitic acids (D-PA) for 6 hours, compared to the signals from HeLa cells grown in DMEM with 1% FBS and 70% D_2_O for 24 hours. The number of lipid droplets formed in the three conditions were quantified (mean ± SD; N > 8). CH_*L*_ signal for D_2_O treatment confirmed the presence of few lipid droplets. (B) SRS signal intensity ratios of third stage larva grown from eggs on 20% D_2_O plates seeded with live or UV-killed *E. coli* OP50 24 hours before imaging. Live bacteria were labeled by deuterium, and killed bacteria was not labeled. (C) Normal fourth stage larva (L4) or dauer larva were transferred from regular NGM plates made of H_2_O to plates made of 20% D_2_O and then imaged at different time points. *E. coli* OP50 were seeded onto the D_2_O plates 24 hours before the experiments. Scale bar = 20 μm. (D) Quantification of the mean intensity over the entire worm body. (E) L4 animals were first grown on D_2_O plates for 1 hour and then transferred to H_2_O plates and imaged at different time points after the transfer. CD_*L*_ mean intensity was plotted. For all *C. elegans* experiments, at least 8 animals were imaged for each condition to calculate mean intensity; mean ± SD of the mean intensity was shown. (F) Ear skin were harvested from adult mice drinking 25% D_2_O for 2, 8, or 26 days, and the sebaceous glands were imaged from the CD_*L*_ and CH_*L*_ channels, from which signals were color-coded in green and red, respectively. (G) Internal capsule of the mouse brain from P5 (5 days postnatal) and P11 pups and adults were sectioned and imaged. Pups were fed on milk produced by mother mice drinking 25% D_2_O for 6 days before imaging, and adults drank 25% D_2_O for 9 days before imaging. (H) White and brown adipose tissues from juvenile (P25) and adult (3-month old) mice drinking 25% D_2_O for 5 days were imaged. Difference between WAT (‘W’) and BAT (‘B’) were shown, as an example, by the enlarged regions (dashed square) of the adult tissues; fluorescence signal excited at 488 nm and collected at 525 nm were shown in purple. (I) Adipose tissues from wild-type and *ob/ob* adult mice that drank 25% D_2_O for 24 days. Scale bar = 20 μm. (J-K) Quantification of data in (C) and (D). For mouse experiments, at least three mice were used for each condition and multiple fields were imaged for each tissue. Mean ± SD was shown.

The large number of lipid droplets caused by the treatment of D-FAs even at low concentrations (10 μM) suggests that exogenous fatty acids likely perturbed native cellular metabolism. Moreover, different types of fatty acids altered lipid metabolism in distinct ways; for the same HeLa cells, treatment of 10 μM D-OA generated remarkably more lipid droplets than D-PA (Figure 4A). Our earlier study also found that PA but not the unsaturated OA, when applied at higher concentration (∼200 μM), drove the formation of solid-like microdomain and membrane phase separation in endoplasmic reticulum^32^. Thus, compared to D-FAs, D_2_O, as a metabolic bystander, would be a much better probe for monitoring endogenous lipogenesis in general.

### DO-SRS tracks lipid metabolism in *C. elegans*

In animals, we first applied DO-SRS imaging to assess how much *de novo* synthesis contributes to the production of total lipids in *C. elegans*, which relies on both dietary uptake (from food source *E. coli* bacteria) and *de novo* lipogenesis for total lipid synthesis. For the quantification purpose, we developed CD_*L*_/CH_*L*_ as a ratiometric indicator for the amount of newly synthesized lipids normalized against variations among individuals and local heterogeneity within the same tissue (see Methods). We found that, when both growing on 20% D_2_O plates, animals fed on non-labeled bacteria (grown in H_2_O) had much lower CD_*L*_/CH_*L*_ ratio (∼3.1%) than animals fed on D-labeled bacteria (∼22.3%; Figure 4B and Supplementary Figure 3A), indicating that ∼14% of total lipids were synthesized *de novo* and the rest ∼86% were incorporated or modified from *E. coli* fatty acids. This result agrees with previous mass spectrometry studies, which used dietary ^13^C labeling and found that *C. elegans* only *de novo* synthesized ∼7% of palmitate and 12%∼19% of eighteen-carbon fatty acids from acetyl-CoA, and the remaining were from dietary uptake^34^. However, compared to mass spectrometry, which has ionization and fragmentation bias towards different analytes, DO-SRS offers easier interpretation and better quantification for total lipid signal. Moreover, mass spectrometry method can only measure metabolic incorporation of isotopes in a bulk of thousands of worms, whereas our *in situ* imaging method detects metabolic activity at the individual animal level, requires much less materials, and can reveal variations among individuals.

D_2_O is a better probe than D-FAs to analyze lipogenesis in *C. elegans* because of the following reasons. First, we directly compared CD_*L*_ signals generated by D_2_O probing and D-FA supplementation (Supplementary Figure 3B and C) and found that although both labeled lipid droplets in similar patterns, 20% D_2_O treatment produced over two-fold stronger CD_*L*_ signal than 4 mM deuterated palmitic acid, the highest concentration used in previous studies^31, 33^. Second, D_2_O is able to track the ∼14% lipids generated by *de novo* lipogenesis, whereas D-FA cannot. Third, because bacteria grown on D_2_O plates produce a variety of D-labeled fatty acids, lipids, as well as their metabolic intermediate, which all become dietary nutrient for *C. elegans*, D_2_O probing (by generating D-labeled bacteria) can monitor lipid synthesis more accurately and more robustly than the supplementation of a single type of D-FA. Fourth, D_2_O probing also generates D-labeled protein signals in addition to lipid signals, whereas D-FA treatment does not (Supplementary Figure 3B).

DO-SRS allowed us to study the dynamics of lipid metabolism in *C. elegans*. When transferred from H_2_O plates to 20% D_2_O plates, normal fourth stage larva (L4) showed newly synthesized lipid droplets in the intestine in as early as 20 minutes; CD_*L*_ signal continued to increase for 6 hours and expanded into hypodermis, suggesting fast and robust lipogenesis (Figure 4C). In contrast, when developmentally arrested dauer larva were placed onto D_2_O plates with food (*E. coli* bacteria), new lipid signals did not appear until 6 hours after the transfer and took 16 hours to reach the 6-hour intensity of normal larva (Figure 4D). The different dynamics of normal and dauer larva reflects the additional time required for dauers to exit the non-feeding, diapause stage and to resume metabolic activities and life cycle^35, 36^. Interestingly, with the unprecedented access to the metabolic status of individual dauers, our data connects metabolic dynamics to previously reported changes in transcriptome during this dauer recovery process, because the onset of lipogenic activities closely follows the expression of genes involved in glycolysis, tricarboxylic acid cycle, fatty acid oxidation, and oxidative phosphorylation^36^. This result suggested that the buildup of storage fat in recovering dauers began immediately after the production of ATP from food digestion and may be important for the preparation of dauer-to-L4 molt, during which food intake stopped and CD_*L*_ signal stayed flat (8 to 12 h; Figure 4D).

We can also monitor lipid degradation through a pulse-chase experiment. For example, L4 animals were first pulsed for one hour on 20% D_2_O plates and then transferred back to regular H_2_O plates. CD_*L*_ signal dropped by ∼60% in the first five hours (Figure 4E), suggesting a quite fast lipid turnover in the developing larva. Thus, DO-SRS enabled the visualization of both lipid anabolism and catabolism.

### Visualizing *de novo* lipogenesis and metabolic homeostasis in mice

By adding D_2_O to the drinking water of mice, previous studies showed the incorporation of deuterium into the total biomass in various mouse organs but could not image and differentiate D-labeled lipids and proteins *in situ*^5-8^. Using DO-SRS imaging with macromolecular selectivity, we found that different mouse organs show metabolic preference to either protein or lipid biosynthesis, which reflects their different functions (Supplementary Figure 4A). We first focused on lipid-rich tissues such as sebaceous glands, myelin sheath in the brain, and adipose tissues, which showed strong CD_*L*_ signals and weak CD_*P*_ signal (Figure 4F-I and Supplementary Figure 4A). Because the visualization of *in situ* lipid synthesis dynamics in mouse models had been particularly challenging with previous methods, which mostly capture the static level^37^, here we demonstrate the utility of DO-SRS in studying the development and metabolic homeostasis of the lipid-rich tissues. From those studies, we also gained new insights into mammalian lipogenesis.

First, we analyzed the spatiotemporal dynamics of lipogenic activities during holocrine secretion. Previous studies imaged the total lipids in sebaceous glands using label-free coherent anti-Stokes Raman scattering (CARS) microscopy and found that sebocytes near the duct had the highest abundance of lipids^38^. But, whether those cells also have the highest lipogenic activity is unclear. By imaging sebaceous glands collected under the ear skin of 3-month old mice that drank 25% D_2_O for 2, 8, or 26 days, respectively, we found that active lipid synthesis occurred mostly in the peripheral sebocytes at day 2, although CH_*L*_ images indicated the presence of a large amount of lipids throughout the entire gland (Figure 4F). The CD_*L*_ signal increased and expanded into the center of the gland later at day 8 and 26, as sebocytes migrated towards the duct and accumulated more lipids (Figure 4F and Supplementary Figure 3D). These results suggest that the outermost early sebocytes, instead of the inner mature sebocytes, had the highest lipogenic activity. At day 26, CD_*L*_/CH_*L*_ ratio reached ∼0.2, which is close to the maximal D:H ratio (17.6∼21.2%) body water from 15∼17% D_2_O enrichment; thus, all the observed lipids were newly synthesized and a likely complete lipid turn-over had occurred.

Second, we visualized the myelination dynamics of axon bundles in the developing brain. Previous studies indicated that myelinogenesis occurs predominantly postnatally in mammals^39^, but it is rather difficult to obtain the precise timing of myelination in a specific region of the brain with traditional methods. By feeding pups for 6 days (P0-P5 or P6-P11) with milk from mother that drank 25% D_2_O, we observed strong bundle-like CD_*L*_ signal labeling the myelinating thalamocortical fibers in the internal capsule in the second postnatal week (imaged at P11) but not in the first week (imaged at P5) and not in adults (Figure 4G). Interestingly, previous studies observed organized growth of thalamocortical projection during the first postnatal week^40^, and our data suggests myelination of those fibers occurred shortly after the axonal development within a stringent time window. In contrast, other axonal fiber-rich structures like hippocampus and olfactory bulbs, which mostly developed embryonically, did not show significant change in myelination activities during the same period of time (Figure 4G and Supplementary Figure 3E).

Third, we observed different metabolic dynamics of brown adipose tissue (BAT) and white adipose tissue (WAT) *in situ* during development and disease. BAT and WAT were identified based on 1) that adipocytes in WAT contain one single, large fat droplet, and adipocytes in BAT contain many small lipid droplets; and 2) that BAT have stronger autofluorescence than WAT due to high levels of cellular NADH and flavin. We found that BAT has higher lipogenic rate (CD_*L*_/CH_*L*_ ratio) in juvenile (P25) mice than in adult mice (Figure 4H and J), consistent with the important thermogenic function of BAT in young animals^41^. WAT serves as energy storage; and age-related increase in the percentage of body fat is often attributed to decrease in resting metabolic rate^42^. However, we found that WAT in adult mice synthesizes and accumulates more fat than WAT in juveniles within the same D_2_O probing period, suggesting that increased lipogenesis is also responsible for fat accumulation in adults. Genetically obese (leptin-deficient *ob/ob*) mice also showed much higher lipogenic activities in WAT than did the wild-type mice, indicating that obesity induces ectopic lipogenesis, in addition to fat accumulation (Figure 4I and K). Hence, DO-SRS can not only track the metabolic abnormality of lipid disorders in animal models but also can serve as a general phenotyping tool for fat metabolism and energy homeostasis.

### DO-SRS enables *in vivo* tracking of *de novo* protein synthesis

Protein synthesis activity is another major component of metabolic dynamics. To visualize newly synthesized proteins in mice *in vivo*, previous studies administered 2.5 mg/ml of D-labeled amino acids (D-AAs) *via* drinking water for 12 days and observed CD signals only in liver and intestine tissues but no other organs^43^. This tissue bias may be explained by the unequal uptake of orally administered D-AAs among different organs, heterogeneity of AA pools, and the poor labeling efficiency of D-AAs^28^. Moreover, the possible varying incompleteness of mixing D-AAs with pre-existing AA pool also makes the measurement of protein synthesis rate complicated. In contrast, D_2_O would have higher labeling efficiency and consistency because D_2_O freely diffuses into all tissues and rapidly labels free AAs through transamination^28^, making the CD_*P*_ signal generated by D_2_O probing more accurately reflect the distribution of newly synthesized proteins than the signal from D-AAs labeling.

We note that the D-AA concentration (or density) in the free AA pool inside cells (<10 mM)^44^ is below our detection limit (∼20 mM for singly deuterated molecules), so CD_*P*_ signal only reports D-AAs incorporated into proteins. In addition, Busch *et al.*^28^ found that there was no post-translational labelling of proteins by D_2_O-derived deuterium, thus D_2_O probing only tracks protein synthesis and not post-translational modification.

To demonstrate that D_2_O is an efficient, consistent, and cost-effective tracer of *de novo* protein synthesis, we performed a direct comparison between 8-day administration of 25% D_2_O and 2 mg/ml of D-AAs both *via* drinking water. Indeed, D-AAs treatment only produced significant CD_*P*_ signals in the digestive tract and liver, where D_2_O probing resulted in much stronger signal (Figure 5). In the pancreas, D-AAs treatment generated weak and sparse signal for protein synthesis activity, whereas D_2_O probing was much more sensitive and revealed broader regions of the tissue that actively synthesized proteins. Moreover, in organs, such as hippocampus, cortex, muscle, and thymus, oral intake of D-AAs hardly produced any CD_*P*_ signal due to tissue bias, whereas D_2_O probing was not subjected to such bias and generated strong CD_*P*_ signal labeling newly synthesized proteins. In terms of the cost, D_2_O administration (∼$0.67/mouse/day at 25%) is 15 times cheaper than D-AAs treatment (∼$10/mouse/day at 2 mg/ml), making D_2_O a much more economical tracer for long-term *in vivo* labeling of slow-turnover proteins.

**Figure 5.**
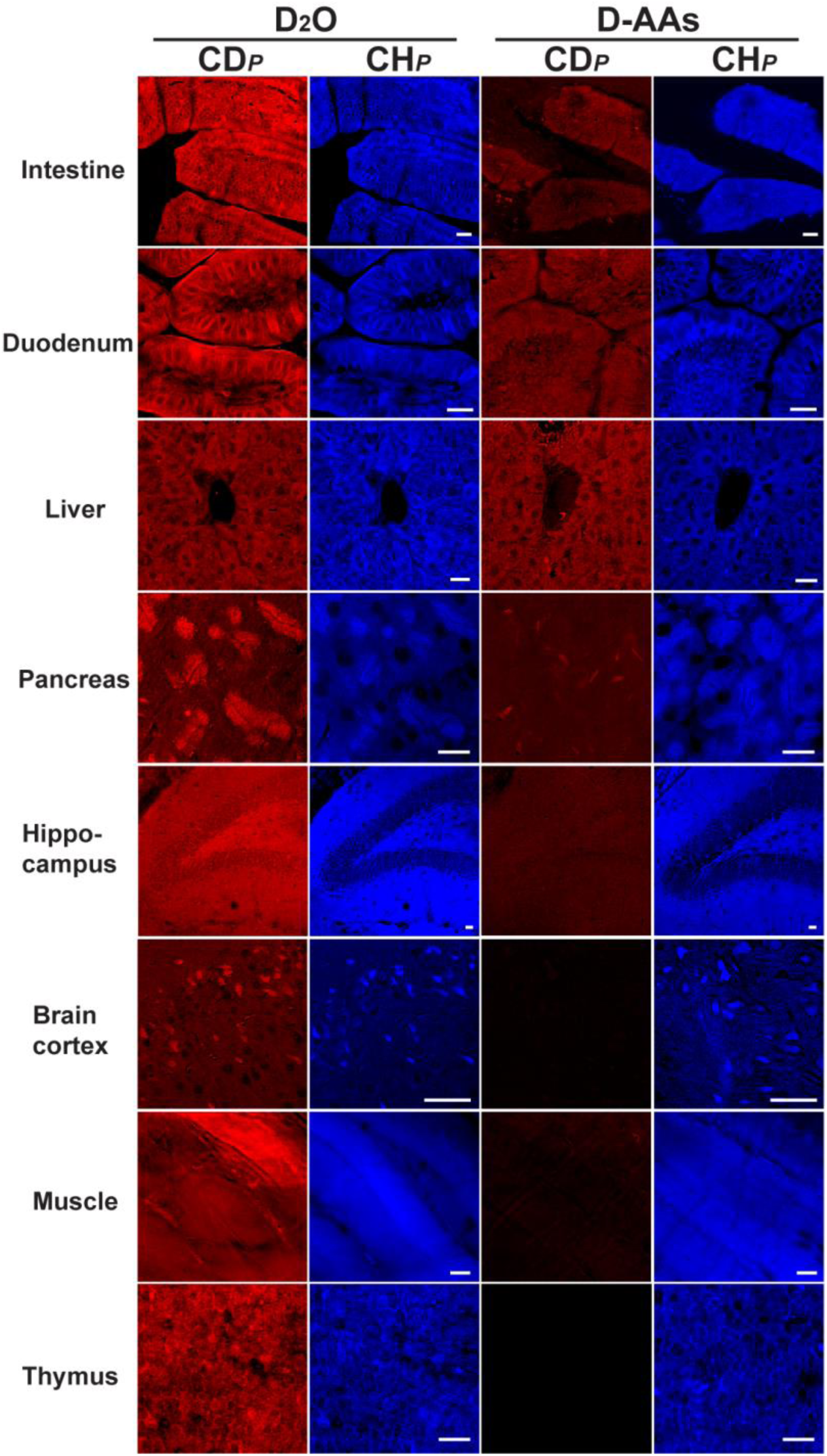
DO-SRS enables tracking of *in vivo* protein synthesis. A variety of protein-rich organs were collected from adult mice that were administrated with 25% D_2_O or 2 mg/ml of D-labeled amino acids (D-AAs) in drinking water for 8 days and then imaged for CD_*P*_ and CH_*P*_ signals.

Although non-drinking administration such as injection of D-AAs directly into the bloodstream *via* carotid artery could enhance CD signals and allow the labeling of proteins in organs, such as pancreas and brain cortex, we found that the signal is still weaker than that generated by D_2_O probing. In general, D-AAs injection to bloodstream only labels tissues with strong protein synthesis activity and is less sensitive than D_2_O treatment, which can reveal low levels of *de novo* protein biosynthesis activities. Furthermore, for organs like hippocampus, injection of D-AAs still could not generate any clear CD_*P*_ signal (Supplementary Figure 4B), suggesting that the D-AA labeling bias against certain tissues is probably inherent to the probe and could not be overcome by increasing D-AA concentration. Compared to the infusion of D-AAs, D_2_O administration through drinking water is also much more convenient, enabling large-scale animal experiments.

### Simultaneous visualization of lipid and protein metabolism *in situ* reveals their different spatial and temporal dynamics

Another major advantage of DO-SRS, compared to D-FA or D-AA treatment alone, is the ability to simultaneously acquire signals for D-labeled lipids and proteins and then resolve their accurate distribution on the same sample through spectral unmixing, with only one probe (D_2_O). This unprecedented capacity allows the study of both lipid and protein metabolism at the same time in an integrated manner.

We first visualized protein and lipid synthesis in *C. elegans* simultaneously during germline development and revealed their different metabolic dynamics. Previous studies suggested that cholesterol, fatty acids, and other nutrients are transported to developing oocytes *via* yolk particles, but it is unclear when exactly lipid deposition occurs during germ cell development and whether there is any difference between protein and lipid accumulation in oocytes^45^. We found that the mitotic and meiotic germ cells showed very active protein synthesis (CD_*P*_) but low level of lipid synthesis (CD_*L*_) signals in L4 animals, whereas the post-pachytene, maturing oocytes in day 0 adults accumulated significant amount of newly synthesized lipids in a 3-hour period (Figure 6A and B). Thus, our results suggest continuous protein synthesis and accumulation throughout germline development and temporally restricted lipid deposition into late-stage oocytes.

**Figure 6.**
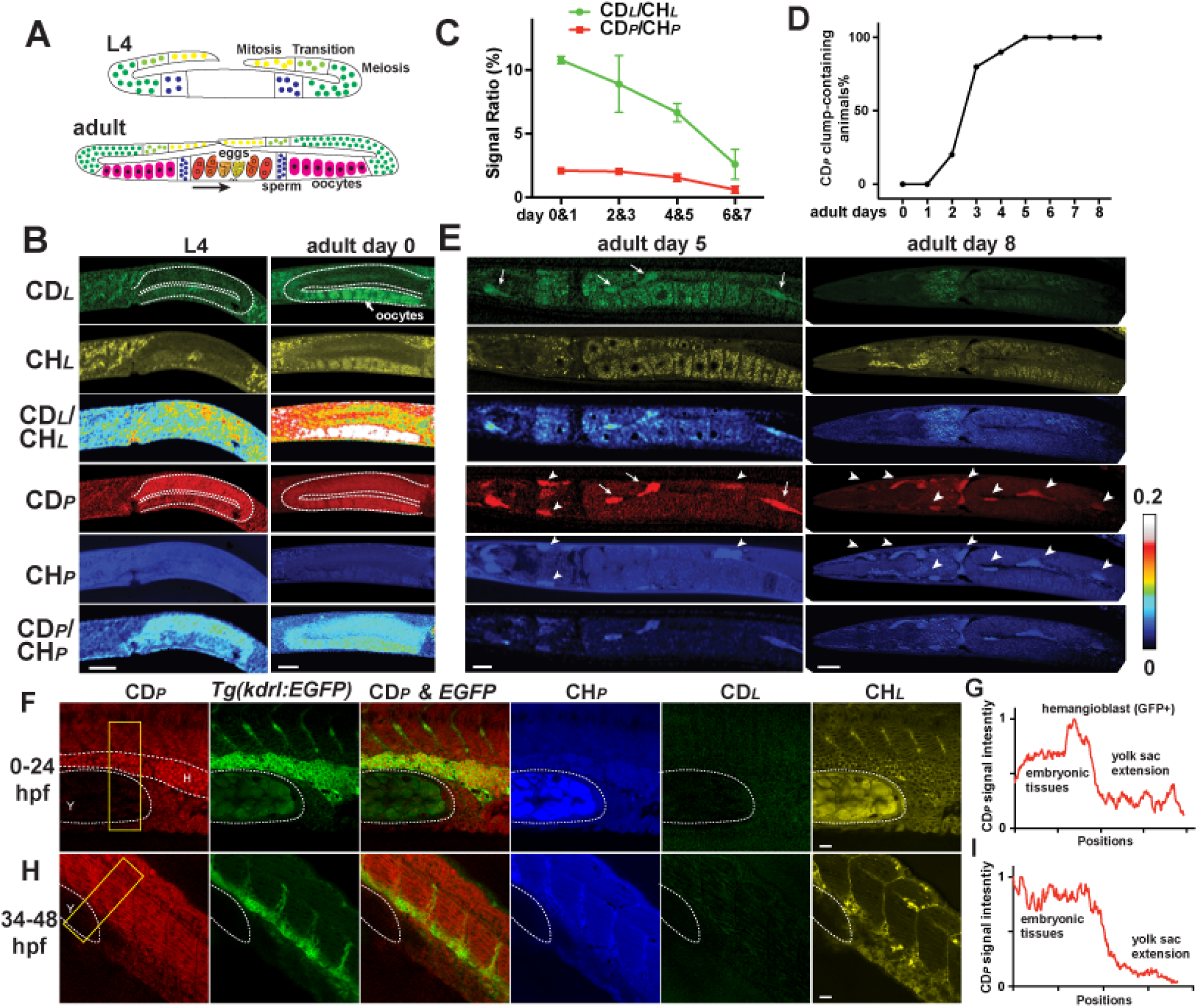
DO-SRS visualizes *in vivo* protein and lipid metabolism simultaneously. (A) Cartoons depict germline development^61^ with the following color scheme: yellow mitotic region, light green transition (early prophase of meiosis I), dark green pachytene, dark blue spermatogenesis, and pink oogenesis. In adults, a color gradient from orange to dark yellow indicate the development (arrow) from newly fertilized eggs to 32-cell embryo, which would be expelled *via* the vulva (triangles). (B) L4 animals and day 0 adults were grown on 20% D_2_O plates for 3 hours before imaging (dashed outline indicates the gonad). Scale bar = 20 μm. (C) Day 0-7 adults were grown on 20% D_2_O plates for 3 hours and then imaged. Mean intensity ratios (mean ± SD) were shown for animals divided into 4 age groups. N > 10 in each group. (D) Day 0 to day 8 adults were transferred from regular NGM plates to 20% D_2_O plates and then imaged in 3 hours. The percentage of animals that showed CD signals in clumps in the body cavity and outside of the oocytes were shown; N > 6 for each time point. (E) SRS images of day 5 and day 8 adults after 3-hour D_2_O probing. Arrows indicate newly formed lipid and protein accumulations only labeled by CD signals, whereas arrowheads indicate pre-existing mass labeled by CH signals. (F-I) SRS signal and the colocalization with fluorescence from *Tg(kdrl::EGFP)* reporter in zebrafish embryos that were incubated in egg solution containing 20% D_2_O from 0 to 24 hpf or from 34 to 48 hpf. Dashed curves outline the yolk sac extension (Y) and the GFP-positive hemangioblast (H) that has strong CD_*P*_ signal. (J) and (L) show intensity profiles that quantifies the CD_*P*_ signal within the yellow rectangle for 0-24 and 34-48 hpf probing, respectively. X axis shows the position along the length of the box from top to bottom, and Y axis shows the average intensity across the width of the box.

Lipid and protein metabolism also showed different age-related changes in *C. elegans*. In general, overall lipid synthesis rate dropped continuously from adult day 0 to day 7, indicating a decline in lipogenesis during aging. The rate of protein synthesis declined most significantly after adult day 5, showing different dynamics from lipids (Figure 6C).

The subcellular resolution afforded by DO-SRS also allowed us to reveal distinct spatial patterns of biosynthesis. Previous studies reported unregulated synthesis and body-wide accumulation of yolk proteins and lipids in the absence of oogenesis in post-reproductive *C. elegans* using either yolk protein::GFP reporter or electron microscopy^46, 47^. However, fluorescent labeling has high background and cannot differentiate proteins from lipids, and EM studies revealed yolk proteins and lipids as different electron-dense materials but are laborious and time-consuming. Importantly, both methods provide only a static view of yolk production, and it is not clear whether the pattern of yolk synthesis changes as the adults age.

Using DO-SRS, we found that from adult day 3 the majority of worms showed significant CD_*L*_ and CD_*P*_ signal as clumps in the intestine and throughout the body cavity after a 3-hour D_2_O probing (Figure 6D and E). CD_*P*_ signal appeared to be stronger than CD_*L*_ signal in older adults (e.g. day 8) but not in younger adults (e.g. day 5), indicating more persistent yolk protein synthesis compared to lipogenesis. Our images not only provided a direct visualization of yolk lipid and protein production but also surprisingly revealed some spatial restriction for the previously considered unregulated biosynthesis of yolk materials. For example, we found that CD_*P*_ signals appeared in both pre-existing mass and newly formed clumps in younger adults (e.g. day 5), suggesting that newly synthesized proteins both accumulated into existing aggregates and formed new aggregates. However, CD_*P*_ signals in older adults (e.g. day 8) only emerged in pre-existing mass, indicating that newly made proteins were only deposited into pre-formed aggregates in aged animals. Thus, our observation revealed an aging-dependent aggregation pattern for newly synthesized yolk proteins. DO-SRS may serve as a useful tool to study protein aggregation *in situ*.

Applying this method to zebrafish, we also identified difference in protein and lipid metabolism during embryonic development. By incubating the embryos in egg water containing 20% D_2_O for 24 hours, we observed significant CD_*P*_ signal and little CD_*L*_ signal in all embryonically derived tissues, which suggests that the rate of protein synthesis, in general, is much higher than the rate of lipogenesis during embryonic cell division. The zebrafish yolk sac, however, showed very weak signal for both CD_*P*_ and CD_*L*_ channel despite significant CH_*L*_ and CH_*P*_ signals from maternally deposited lipids and proteins (Figure 6F). Since the yolk and blastoderm had similar water permeability^48^, our result suggests very little zygotic biosynthesis in the yolk independently of maternal contribution. Overall, our observations relied on the *in situ* separation of D-labeled proteins and lipids, which cannot be easily achieved by previous methods.

We also used DO-SRS in conjunction with fluorescent microscopy to track the metabolic activity of specific cell lineages during development. Probing the embryos from zygote to 24 hours post-fertilization (hpf), we found that a group of hemangioblast cells labeled by *Tg(kdrl:EGFP)*^49^ demonstrated the strongest CD_*P*_ signal (Figure 6F and G). Those cells originate from lateral plate mesoderm and give rise to endothelial and hematopoietic lineages^50^; their strong CD_*P*_ signal indicates active protein biosynthesis, possibly due to fast proliferation, during the 0-24 hpf period. D_2_O probing of later embryonic stages, 24-34 and 34-48 hpf, found that the *EGFP+* cells, which labeled mostly differentiated endothelial cells then^51^, no longer showed stronger CD_*P*_ signal than the surrounding tissues (Figure 6H and I and Supplementary Figure 4). Thus, by co-labeling with lineage-specific fluorescent marker, we can track time-dependent metabolic activities of particular cell types during lineage progression.

### DO-SRS visualizes tumor boundary and metabolic heterogeneity

An immediate biomedical application of DO-SRS is to visualize tumor boundaries and intratumoral metabolic heterogeneity. Although label-free SRS identified the boundary between glioblastoma and normal brain tissues^52^, it relies on the protein/lipid compositional difference between the tumor and normal tissues—brain areas had higher myelin-derived lipid concentration than tumors, which may not apply to other type of tumors. In contrast, DO-SRS can reveal tumor boundaries by tumor’s inherently higher metabolic activities than the surrounding normal tissues. For example, giving glioblastoma-bearing mice 25% D_2_O for 15 days, we observed both stronger CD_*L*_ and CD_*P*_ signals in the tumor tissue compared to the nearby brain tissues, even though the brain region had high total lipids in the CH_*L*_ channel (Figure 7A). Unlike the brain tumors, in the subcutaneous xenograft of colon tumor, the tumor and the surrounding skin tissue had similar composition of proteins and lipids and, hence, were indistinguishable by label-free SRS from their CH_*L*_ and CH_*P*_ signal. However, through D_2_O probing, the tumor showed higher level of lipogenesis than the skin and became readily identifiable in the CD_*L*_ channel (Figure 7B). Thus, this example showcases that DO-SRS could be a more general and applicable method of detecting tumor boundaries than the label-free SRS.

**Figure 7.**
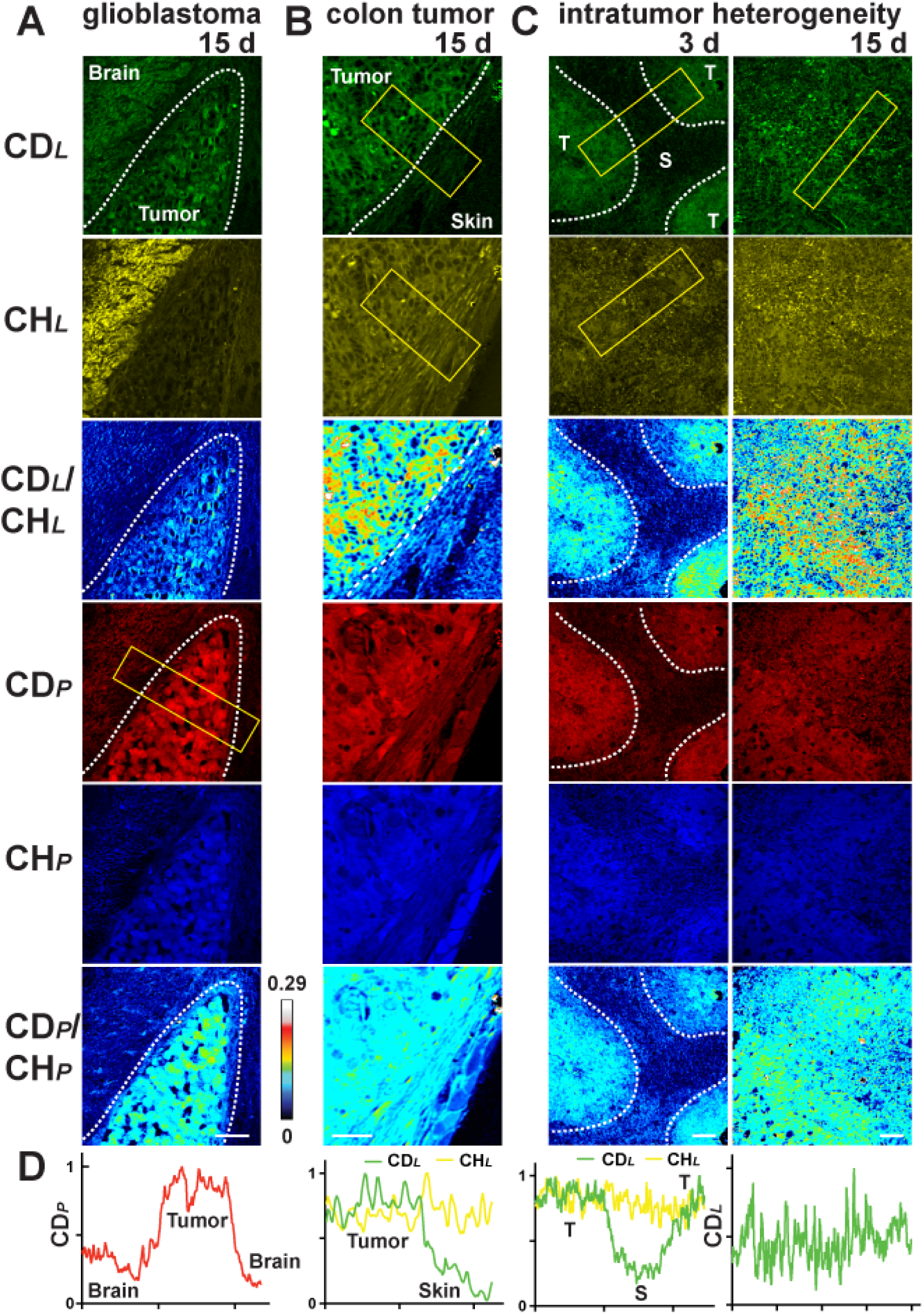
DO-SRS identifies tumor boundaries and metabolic heterogeneity. (A) Intracranial xenograft glioblastoma in mouse brain were excised, sectioned, and imaged after the tumor-bearing mice drank 25% D_2_O for 15 days. Color scheme is similar to that in Figure 2. Dashed curves highlight the tumor-brain boundary visualized by CD_*P*_ signal. Intensity profile quantifies the CD_*P*_ signal within the yellow rectangle with X axis showing the position along the length of the box and Y axis showing the average intensity across the width of the box. (B) Methods similar to (A) was used to visualize the tumor-skin boundary of subcutaneously xenografted colon tumor from the CD_*L*_ channel. (C) The interior of the colon tumor xenografts was imaged after the tumor-bearing mice drank 25% D_2_O for 3 or 15 days. Dashed lines indicate the boundary between the tumor cells (T) and the recruited stromal cells (S), which were identified by the CD_*L*_ signals at 3 days. Tumor and stromal cells were also identified by their different morphologies. Scale bar = 20 μm. (D) Intensity profile quantifies the CD_*P*_ or CD_*L*_ signal within the yellow rectangular with X axis showing the position along the length of the box and Y axis showing the average intensity across the width of the box.

Intratumoral metabolic heterogeneity is considered a driver of tumor aggressiveness and has been under intensive study due to its fundamental importance as well as prognostic significance^53^. In the colon tumor xenograft, cancer cells recruited stromal cells from the nearby normal tissues and developed into solid tumor. 3-day D_2_O probing revealed that human tumor cells had stronger biogenesis for both proteins and lipids than the recruited mouse stroma, showing metabolic heterogeneity inside the solid tumor (Figure 7C). Interestingly, the difference between tumor and stroma became less pronounced after 15-day D_2_O intake, suggesting that the stromal cells also had significant albeit slower metabolic activity presumably to support tumor growth (Figure 7D). Therefore, DO-SRS can visualize the internal structure and metabolism of solid tumors with cellular resolution.

## Discussion

In summary, we developed and demonstrated DO-SRS as a nondestructive, noninvasive, and background-free imaging method that can be used to visualize metabolic dynamics of proteins, lipids, and DNA in a variety of model organisms. DO-SRS allows for *in situ* visualization of *de novo* lipogenesis and protein synthesis in animals at an unprecedentedly low cost and without tissue bias, representing important technical advance. In particular, the ability to simultaneously image newly synthesized lipids and proteins allowed us to gain new insights into the metabolic basis of several biological processes, such as the distinct protein and lipid synthesis patterns in *C. elegans* germline and the prevalence of protein biosynthesis and the lack of lipogenesis in zebrafish embryos. We also showcased the tremendous potential of using the unmixed CD_*L*_ and CD_*P*_ signals to identify tumor boundaries and to detect intratumoral heterogeneity. Therefore, DO-SRS serves as a powerful new tool to facilitate the study of metabolic activity-dependent biological processes.

Several mass spectrometry imaging (MSI) methods have been developed to capture the spatial distribution of D-labeled biomolecules in fixed tissues with chemical resolvability, mostly among low-molecular-weight metabolite and peptides^54-56^. Although the high molecular specificity and sensitivity of MSI methods cannot be achieved by DO-SRS currently, it still serves as an important complementary approach to MSI. All MSI techniques are essentially destructive surface analysis, require intensive computation, and have bias towards easily ionized and desorbed analytes. DO-SRS, on the contrary, provides straightforward and quantitative interpretation of metabolic activities with macromolecular selectivity in three-dimensional living tissues.

Using devices similar to the coherent Raman scattering endoscopes^57^, we envision that DO-SRS, with its ability to image D-labeled macromolecules in living animals (Figure 1D), could be applied to visualize metabolic patterns of internal organs, cortical metabolism for brain activities, and tumor metabolism for cancer progression through optical biopsy. Importantly, the sensitivity of this method is high enough to operate in the range of low D_2_O enrichment that is safe for humans. Given that SRS imaging has been demonstrated in humans before^58^ and the recent development of high-speed, volumetric stimulated Raman projection (SRP) microscopy and tomography offers promise in deep-tissue, large-volume, *in vivo* imaging^59^, we expect DO-SRS, as a safe and non-invasive imaging technique, to have clinical application in tracking metabolic activities in humans.

## Acknowledgement

We thank Dr. Martin Chalfie for critical comments on the manuscript. C. Zheng is supported by NIGMS grants (GM30997 and GM122522) to Martin Chalfie. C.D.S. Tomas and K. Targoff are supported by NHLBI R01 (HL131438-01A1). W. Min acknowledges support from the NIH Director’s New Innovator Award (1DP2EB016573), R01 (EB020892), the US Army Research Office (W911NF-12-1-0594), the Alfred P. Sloan Foundation, and the Camille and Henry Dreyfus Foundation.

## Author Contributions

L.S., C.Z., Y.S, Z.C, and W.M. conceived the project. L.S., C.Z., Y.S., E.S.S, L.Z., M.W., C.L., and C.D.S.T. performed experiments. C.Z. and L.S. wrote the original manuscript. C.Z., L.S., Y.S., W.M., Z.C., C.D.S.T., and K.T. edited the manuscript. K.T. and W.M secured funding.

## Declare of Interests

The authors declare no competing interests.

## Reference

1. Kim, M.M., Parolia, A., Dunphy, M.P. & Venneti, S. Non-invasive metabolic imaging of brain tumours in the era of precision medicine. Nat Rev Clin Oncol 13, 725–739 (2016).

2. Musat, N., Foster, R., Vagner, T., Adam, B. & Kuypers, M.M. Detecting metabolic activities in single cells, with emphasis on nanoSIMS. FEMS Microbiol Rev 36, 486–511 (2012).

3. Lechene, C. et al. High-resolution quantitative imaging of mammalian and bacterial cells using stable isotope mass spectrometry. J Biol 5, 20 (2006).

4. Steinhauser, M.L. et al. Multi-isotope imaging mass spectrometry quantifies stem cell division and metabolism. Nature 481, 516–519 (2012).

5. Miyagi, M. & Kasumov, T. Monitoring the synthesis of biomolecules using mass spectrometry. Philos Trans A Math Phys Eng Sci 374 (2016).

6. Previs, S.F. et al. New methodologies for studying lipid synthesis and turnover: looking backwards to enable moving forwards. Biochim Biophys Acta 1842, 402–413 (2014).

7. Foletta, V.C. et al. Analysis of Mammalian Cell Proliferation and Macromolecule Synthesis Using Deuterated Water and Gas Chromatography-Mass Spectrometry. Metabolites 6 (2016).

8. Kloehn, J., Saunders, E.C., O’Callaghan, S., Dagley, M.J. & McConville, M.J. Characterization of metabolically quiescent Leishmania parasites in murine lesions using heavy water labeling. PLoS Pathog 11, e1004683 (2015).

9. Berry, D. et al. Tracking heavy water (D2O) incorporation for identifying and sorting active microbial cells. Proc Natl Acad Sci U S A 112, E194–203 (2015).

10. Tao, Y. et al. Metabolic-Activity-Based Assessment of Antimicrobial Effects by D2O-Labeled Single-Cell Raman Microspectroscopy. Anal Chem 89, 4108–4115 (2017).

11. Freudiger, C.W. et al. Label-free biomedical imaging with high sensitivity by stimulated Raman scattering microscopy. Science 322, 1857–1861 (2008).

12. Min, W., Freudiger, C.W., Lu, S. & Xie, X.S. Coherent nonlinear optical imaging: beyond fluorescence microscopy. Annu Rev Phys Chem 62, 507–530 (2011).

13. Cheng, J.X. & Xie, X.S. Vibrational spectroscopic imaging of living systems: An emerging platform for biology and medicine. Science 350, aaa8870 (2015).

14. Wei, L. et al. Live-Cell Bioorthogonal Chemical Imaging: Stimulated Raman Scattering Microscopy of Vibrational Probes. Acc Chem Res 49, 1494–1502 (2016).

15. Wei, L. et al. Super-multiplex vibrational imaging. Nature 544, 465–470 (2017).

16. Camp, C.H. & Cicerone, M.T. Chemically sensitive bioimaging with coherent Raman scattering. Nat Photonics 9, 295–305 (2015).

17. Valencia, M.E., Aleman-Mateo, H., Salazar, G. & Hernandez Triana, M. Body composition by hydrometry (deuterium oxide dilution) and bioelectrical impedance in subjects aged >60 y from rural regions of Cuba, Chile and Mexico. Int J Obes Relat Metab Disord 27, 848–855 (2003).

18. Schoeller, D.A. Recent advances from application of doubly labeled water to measurement of human energy expenditure. J Nutr 129, 1765–1768 (1999).

19. Jones, P.J. & Leatherdale, S.T. Stable isotopes in clinical research: safety reaffirmed. Clin Sci (Lond) 80, 277–280 (1991).

20. Guillermier, C. et al. Imaging mass spectrometry demonstrates age-related decline in human adipose plasticity. JCI Insight 2, e90349 (2017).

21. Neese, R.A. et al. Measurement in vivo of proliferation rates of slow turnover cells by 2H2O labeling of the deoxyribose moiety of DNA. Proc Natl Acad Sci U S A 99, 15345–15350 (2002).

22. Hodel, A., Gebbers, J.O., Cottier, H. & Laissue, J.A. Effects of prolonged moderate body deuteration on proliferative activity in major cell renewal systems in mice. Life Sci 30, 1987–1996 (1982).

23. Peng, S.K., Ho, K.J. & Taylor, C.B. Biologic effects of prolonged exposure to deuterium oxide. A behavioral, metabolic, and morphologic study. Arch Pathol 94, 81–89 (1972).

24. Lu, F.K. et al. Multicolor stimulated Raman scattering (SRS) microscopy. Mol Phys110, 1927–1932 (2012).

25. Yu, Z.L. et al. Label-free chemical imaging in vivo: three-dimensional non-invasive microscopic observation of amphioxus notochord through stimulated Raman scattering (SRS). Chemical Science 3, 2646–2654 (2012).

26. Lu, F.K. et al. Label-free DNA imaging in vivo with stimulated Raman scattering microscopy. Proc Natl Acad Sci U S A 112, 11624–11629 (2015).

27. Diem, M., Polavarapu, P.L., Oboodi, M. & Nafie, L.A. Vibrational circular dichroism in amino acids and peptides. 4. Vibrational analysis, assignments, and solution-phase Raman spectra of deuterated isotopomers of alanine. J. Am. Chem. Soc. 104, 3329–3336 (1985).

28. Busch, R. et al. Measurement of protein turnover rates by heavy water labeling of nonessential amino acids. Biochim Biophys Acta 1760, 730–744 (2006).

29. Lewis, C.A. et al. Tracing compartmentalized NADPH metabolism in the cytosol and mitochondria of mammalian cells. Mol Cell 55, 253–263 (2014).

30. Hu, F., Lamprecht, M.R., Wei, L., Morrison, B. & Min, W. Bioorthogonal chemical imaging of metabolic activities in live mammalian hippocampal tissues with stimulated Raman scattering. Sci Rep 6, 39660 (2016).

31. Fu, D. et al. In vivo metabolic fingerprinting of neutral lipids with hyperspectral stimulated Raman scattering microscopy. J Am Chem Soc 136, 8820–8828 (2014).

32. Shen, Y. et al. Metabolic activity induces membrane phase separation in endoplasmic reticulum. Proc Natl Acad Sci U S A 114, 13394–13399 (2017).

33. Yu, Y., Mutlu, A.S., Liu, H. & Wang, M.C. High-throughput screens using photo-highlighting discover BMP signaling in mitochondrial lipid oxidation. Nat Commun 8, 865 (2017).

34. Perez, C.L. & Van Gilst, M.R. A 13C isotope labeling strategy reveals the influence of insulin signaling on lipogenesis in C. elegans. Cell Metab 8, 266–274 (2008).

35. Cassada, R.C. & Russell, R.L. The dauerlarva, a post-embryonic developmental variant of the nematode Caenorhabditis elegans. Dev Biol 46, 326–342 (1975).

36. Wang, J. & Kim, S.K. Global analysis of dauer gene expression in Caenorhabditis elegans. Development 130, 1621–1634 (2003).

37. Daemen, S., van Zandvoort, M.A., Parekh, S.H. & Hesselink, M.K. Microscopy tools for the investigation of intracellular lipid storage and dynamics. Mol Metab 5, 153–163 (2016).

38. Jung, Y., Tam, J., Jalian, H.R., Anderson, R.R. & Evans, C.L. Longitudinal, 3D in vivo imaging of sebaceous glands by coherent anti-stokes Raman scattering microscopy: normal function and response to cryotherapy. J Invest Dermatol 135, 39–44 (2015).

39. Bercury, K.K. & Macklin, W.B. Dynamics and mechanisms of CNS myelination. Dev Cell 32, 447–458 (2015).

40. Agmon, A., Yang, L.T., O’Dowd, D.K. & Jones, E.G. Organized growth of thalamocortical axons from the deep tier of terminations into layer IV of developing mouse barrel cortex. J Neurosci 13, 5365–5382 (1993).

41. Saely, C.H., Geiger, K. & Drexel, H. Brown versus white adipose tissue: a mini-review. Gerontology 58, 15–23 (2012).

42. St-Onge, M.P. & Gallagher, D. Body composition changes with aging: the cause or the result of alterations in metabolic rate and macronutrient oxidation? Nutrition 26, 152–155 (2010).

43. Wei, L. et al. Imaging complex protein metabolism in live organisms by stimulated Raman scattering microscopy with isotope labeling. ACS Chem Biol 10, 901–908 (2015).

44. Piez, K.A. & Eagle, H. The free amino acid pool of cultured human cells. J Biol Chem231, 533–545 (1958).

45. Branicky, R., Desjardins, D., Liu, J.L. & Hekimi, S. Lipid transport and signaling in Caenorhabditis elegans. Dev Dyn 239, 1365–1377 (2010).

46. Herndon, L.A. et al. Stochastic and genetic factors influence tissue-specific decline in ageing C. elegans. Nature 419, 808–814 (2002).

47. Epstein, J., Himmelhoch, S. & Gershon, D. Studies on ageing in nematodes III. Electronmicroscopical studies on age-associated cellular damage. Mech Ageing Dev 1, 245–255 (1972).

48. Hagedorn, M., Kleinhans, F.W., Artemov, D. & Pilatus, U. Characterization of a major permeability barrier in the zebrafish embryo. Biol Reprod 59, 1240–1250 (1998).

49. Choi, J. et al. FoxH1 negatively modulates flk1 gene expression and vascular formation in zebrafish. Dev Biol 304, 735–744 (2007).

50. Bertrand, J.Y. & Traver, D. Hematopoietic cell development in the zebrafish embryo. Curr Opin Hematol 16, 243–248 (2009).

51. Jin, S.W., Beis, D., Mitchell, T., Chen, J.N. & Stainier, D.Y. Cellular and molecular analyses of vascular tube and lumen formation in zebrafish. Development 132, 5199–5209 (2005).

52. Ji, M. et al. Rapid, label-free detection of brain tumors with stimulated Raman scattering microscopy. Sci Transl Med 5, 201ra119 (2013).

53. Sengupta, D. & Pratx, G. Imaging metabolic heterogeneity in cancer. Mol Cancer 15, 4 (2016).

54. Steinhauser, M.L., Guillermier, C., Wang, M. & Lechene, C.P. Quantifying cell division with deuterated water and multi-isotope imaging mass spectrometry (MIMS). Surf Interface Anal 46, 161–164 (2014).

55. Louie, K.B. et al. Mass spectrometry imaging for in situ kinetic histochemistry. Sci Rep 3, 1656 (2013).

56. Gemperline, E., Chen, B. & Li, L. Challenges and recent advances in mass spectrometric imaging of neurotransmitters. Bioanalysis 6, 525–540 (2014).

57. Saar, B.G., Johnston, R.S., Freudiger, C.W., Xie, X.S. & Seibel, E.J. Coherent Raman scanning fiber endoscopy. Opt Lett 36, 2396–2398 (2011).

58. Saar, B.G. et al. Video-rate molecular imaging in vivo with stimulated Raman scattering. Science 330, 1368–1370 (2010).

59. Chen, X. et al. Volumetric chemical imaging by stimulated Raman projection microscopy and tomography. Nat Commun 8, 15117 (2017).

60. Ujike, T. & Tominaga, Y. Raman spectral analysis of liquid ammonia and aqueous solution of ammonia. J Raman Spectrosc 33, 485–493 (2002).

61. Hubbard, E.J. & Greenstein, D. Introduction to the germ line. WormBook, 1-4 (2005).

