## Supplementary Materials for "Optical Imaging of Metabolic Dynamics in Animals"

### Materials and Methods

#### Raman Spectroscopy

##### Spontaneous Raman spectroscopy

Spontaneous Raman scattering spectra are acquired as previously described[1] on an upright confocal Raman microspectrometer (Xplora, Horiba Jobin Yvon) with 532 nm diode laser source and 1800 l/mm grating at room temperature. The excitation power is ~40 mW after passing through a 50× air objective (MPlan N, 0.75 N.A., Olympus), and 40s acquisition time accumulated by 22 was used to collect Raman spectra of all samples at a single point under identical conditions. For cultured cells, The Raman background of water and cover glass is removed from all cell spectra by subtracting the signal at empty space from the signals collected from cells.

##### Stimulated Raman Scattering Microscopy

Similar to a previously described setup[2], we used an inverted laser-scanning microscope (FV1200 MPE, Olympus) optimized for near-IR throughput and a 25× water objective (XLPlan N, 1.05 N.A., MP, Olympus) with high near-IR transmission for SRS imaging. A picoEMERALD system (Applied Physics & Electronics) supplied synchronized pulse pump beam (with tunable 720-990 nm wavelength, 5-6 ps pulse width, and 80-MHz repetition rate) and Stokes (with fixed wavelength at 1064 nm, 6 ps pulse width, and 80 MHz repetition rate). Stokes is modulated at 8MHz by an electronic optic modulator. Transmission of the forward-going Pump and Stokes beams after passing through the samples was collected by a high N.A. oil condenser (N.A. = 1.4). A high O.D. bandpass filter (890/220, Chroma) is used to block the Stokes beam completely and to transmit the pump beam only onto a large area Si photodiode for the detection of the stimulated Raman loss signal. The output current from the photodiode is terminated, filtered, and demodulated by a lock-in amplifier at 8 MHz to ensure shot-noise-limited detection sensitivity. The demodulated signal is fed into analog channel of FV1200 software FluoView 4.1a (Olympus) to form image during laser scanning.

#### Cell culture and treatments

##### Cell culture and imaging

HeLa, COS-7, and U-87 MG cells were obtained from ATCC (Manassas, VA) and cultured in Dulbecco's modified Eagle's medium (DMEM, Thermo Fisher) supplemented with 10% fetal bovine serum (Thermo Fisher) in a humidified 5% $CO_2$ atmosphere at 37°C. LS174T was also obtained from ATCC and cultured in Eagle's Minimum Essential Medium (EMEM from ATCC). To make cell culture medium containing $D_2O$, we used a mixture of $D_2O$ (Sigma-Aldrich, Cat. 151882) and distilled $H_2O$ to dissolve DMEM powder (Sigma) and then sterilized the medium by filtering.

For live-cell imaging, HeLa and COS-7 cells were seeded onto a glass-bottom dish (MatTek) and grown for 6 to 8 hours in regular DMEM before the medium was changed to DMEM containing 70% $D_2O$. Cells were then grown for different periods of time before being placed onto the stage for SRS imaging. When treating cells with D-labeled fatty acids, they were first reacted with sodium hydroxide and then coupled to bovine serum albumin (BSA) in about 2:1 molar ratio to make 2mM stock solution. Those stock solutions were then added to the medium to reach the final working concentration (10 μM).

**Cellular toxicity assay**

To determine cellular toxicity, HeLa cells and COS-7 cells were first seeded in 96-well plates at 500 per $cm^2$ and grown overnight; the culture was then changed to DMEM containing different concentration (0% to 100%) of $D_2O$ for 48 hours. The viability of cell was accessed using the CellTiter-Glo® Luminescent Cell Viability Assay (Promega) according to the manufacturer's protocol. We used cells cultured in pure DMEM medium as control and wells containing only medium without cells for background luminescence. Viability was calculated using the background-corrected absorbance as follows: Viability (%) = Absorbance of experiment well/Absorbance of control well × 100%. Three replicates were performed.

**Treatment of inhibitors**

For inhibitor treatments, cells were first grown in glass bottom dishes with DMEM made of $H_2O$ overnight, and then the medium was changed to 70% $D_2O$ DMEM containing either 10nM fatty acid synthase inhibitor TVB-3166 (Sigma CAS 1533438-83-3) or 1 μM protein synthesis inhibitor anisomycin (Sigma CAS 22862-76-6). Cells were then cultured for 24 hours, imaged by SRS microscopy, and fixed by 4% paraformaldehyde for spontaneous Raman spectroscopy.

Methanol wash was performed on fixed cells in glass-bottom dishes and tissues with 99.9% methanol (Sigma) for 30 minutes and 24 hours, respectively, followed by phosphate buffered saline (PBS) wash. For proteinase K treatment, tissues were first fixed by paraformaldehyde and then treated with 0.2 mg/ml recombinant proteinase K (Roche) at 37°C for 15 minutes and subsequently washed with PBS.

#### Biochemical extraction of macromolecules

##### Extraction of lipids

HeLa cells were grown in DMEM with 70% $D_2O$ for 24 hours, fixed with 4% paraformaldehyde, washed with PBS in petri dishes, and then harvested into 15-mL falcon tubes. 1.3 mL of chloroform and 2.7 mL of methanol were then added to the cells to extract lipids. After centrifugation at 4000 rpm for 5 minutes, the supernatant was transferred to a clean tube. Subsequently, 1 mL of 50 mM citric acid, 2 mL of water, and 1 mL of chloroform were added to the solution and vigorous shaking was used to mix the content. Liquid phases were then separated by centrifugation at 4000 rpm for 10 minutes and the lower phase containing the lipids were transfer to a clean tube. Caps were kept open to allow the evaporation of solvent and the concentration of lipids.

##### Extraction of proteins

Proteins were extracted using Trizol (catalog # 15596026, Life Technologies) according to the manual provided by the manufacturer.

##### Extraction of DNA

DNA were extracted using DNAzol (catalog # 10503027, Thermo Fisher Scientific) according to the protocol provided by the manufacturer.

#### Animal experiments

##### *C. elegans* maintenance and experiments

*C. elegans* wild type (N2) and mutant strains were maintained at 20 °C as previously described[3]. Different concentration of $D_2O$ were added to replace $H_2O$ in nematode growth media (NGM). *E. coli* OP50 were seeded on $D_2O$-containing NGM plates and grown for 24 hours; we then placed worms at different ages onto those plates for various amount of time. To determine the toxicity of $D_2O$, we prepared eggs using hypochlorite, which made the eggshell porous and permeable to $D_2O$, placed them onto plates containing different concentration of $D_2O$, and counted the number of eggs that could not hatch into normal moving larva. We also placed fourth stage larva onto those plates and counted how many animals became sterile and determined the brood size by counting the total number of hatch larvae from the all the eggs laid by one animal.

To prevent deuterium labeling of the bacteria and to feed worms with non-labeled bacteria on $D_2O$ plates, we killed the OP50 bacteria (grown in $H_2O$) immediately after placing them onto $D_2O$ plates with UV light. Worms grown on dead non-labeled bacteria developed normally and had similar level of $CH_2$ and $CH_3$ signal, compared to the controls fed on bacteria that grew on $D_2O$ plates for at least 24 hours.

To supplement deuterated fatty acids to worms, OP50 bacterial culture was mixed well with 4 mM D31-palmitic acid and then seeded onto NGM plates that contained 100% $H_2O$. As controls, the same OP50 bacteria culture were seeded onto NGM plates that contained 20% $D_2O$. 24 hours later, eggs were placed onto those plates and grew until L4 stage before being imaged.

Regular SRS imaging was done after fixation with 4% formaldehyde for 30 minutes and washing with PBS. Live imaging was done by directly transferring animals from $D_2O$ plates onto an 4% agarose pad and mount them in M9 solution containing 0.1 μm polystyrene beads (Polysciences, Inc.) to reduce mobility. For most *C. elegans* experiments, at least 8 animals were imaged for each condition.

**Metabolic activity tracking in zebrafish embryos**

Zebrafish embryos carrying the fluorescent reporters *Tg(kdrl:EGFP)*[4] were incubated in egg water[5] supplemented with 20% $D_2O$ for 24 or 48 hours from 1-cell stage at 28.5°C. For $D_2O$ probing from 24 to 36 hpf (hours post fertilization) and from 34 to 48 hpf, embryos were transferred from 100% $H_2O$ environment to egg water containing 20% $D_2O$. After the incubation, embryos were manually dechorionated and fixed in 4% PFA overnight at 4 °C. Following fixation, the embryos were transferred to PBS prior to imaging. At least four embryos were imaged for each condition. The zebrafish work followed the IACUC (the Institutional Animal Care and Use Committee at Columbia University)-approved protocol AC-AAAJ7554.

**$D_2O$ probing in mice**

Wild-type 3~4 month old adult C57BL/6J and *ob/ob* (B6.Cg-$Lep^{ob}$/J) mice were both obtained from the Jackson Laboratory and were maintained and bred at Columbia University animal facility. For the *ex vivo* mice tissue experiments, mice that drank 25% $D_2O$ for certain days were anesthetized with isoflurane and sacrificed. Various organs and tissues were harvested, fixed with 4% formaldehyde overnight, and then cut into slices of 120 μm thickness using a vibrating blade microtome (Vibratome, Leica) for SRS imaging. For all mice experiments, three mice were used for each condition, and at least three different fields of each tissue were imaged.

For the *in vivo* SRS imaging of ear skin, mice were kept anesthetized with isoflurane while one ear was gently sandwiched between a cover slip and a glass slide, which were then placed onto the imaging stage with a heating pad that keep the body warm during the imaging session.

**Tumor xenografts in mice**

To establish glioblastoma xenograft, we performed intracranial implantation of U-87 MG human glioma cells in nude mice (J:NU). Briefly, the mouse was anesthetized and positioned in stereotaxic instrument (David Kopf Instruments), and then a small section (2 mm in diameter) of the skull is ground with a dental drill until it became soft and translucent. Subsequently, we injected $1.5 \times 10^5$ U87-MG tumor cells (in 3 μl) injected into the frontal region of the cerebral cortex over the course of 5 minutes using a 1.5 mm glass capillary. After the implantation, mouse head skin is then closed with SILK sutures (Harvard Apparatus). Two weeks later, mice bearing glioblastoma started and were kept on drinking 25% $D_2O$ for 15 days before being sacrificed for tumor imaging. For the colon tumor xenograft, we injected subcutaneously $1 \times 10^7$ human colorectal LS174T cells into the lower flank of nude mouse. Ten days later, tumor bearing mice were treated with 25% $D_2O$ as drinking water for 3 or 15 days before being sacrificed for tumor SRS imaging. The experimental protocols for all the mice studies (AC-AAAQ0496) was approved by IACUC.

##### Feeding and Injection of D-labeled amino acids

To feed mice with D-labeled amino acids (D-AAs), D-AA mixture (Catalog # DLM-6819-1, Cambridge Isotope Laboratory) were dissolved in drinking water at 2 mg/ml, and adult mice drank approximately 4 ml of the water every day for 8 days before their organs were harvested for SRS imaging.

To inject D-AAs into the bloodstream *via* carotid artery, mice were anesthetized and kept warm on a heating pad. Neck skin of the mouse was cleaned with 2% chlorhexidine solution followed by 70% isopropyl alcohol and the planned incision area was infiltrated subcutaneously with a 1:10 dilution of 50/50 lidocaine (1%). A ~2cm incision was made along the midline region of the throat to expose the common carotid artery, which was then tied with suture thread (Prolene 86979 Ethicon) on the side closer to heart to stop the blood flow. The artery was then cut with precision stainless micro-scissor (63041-984, VWR), and a polyurethane based Micro-Renathane catheter tube (MRE033, Braintree Scientific, INC) was carefully cannulated into the opening of carotid artery [6]. A syringe filled with 80 mg/ml D-labeled amino acids dissolved in mammalian Ringer solution was then connected with the cannulated tubing, and liquid flow was controlled by a syringe pump (AL-1000, WPI) to inject the solution at 0.01 ml/min perfusion rate for half an hour each session for 2.5 days with a time interval of 2 hours.

#### Spectral Unmixing

##### $CD_L/CD_P/CD_{DNA}$ three-component unmixing

Three-component unmixing of $CH_L/CH_P/CH_{DNA}$ signals were previously reported[7]. In this study, we developed a similar method to unmix CD signals. Briefly, we first acquired SRS signals from 2135 $cm^{-1}$, 2185 $cm^{-1}$, and 2210 $cm^{-1}$ bearing the intrinsic features of lipids, proteins, and DNA, respectively, and then used a linear combination of the three signals with coefficients to determine the amount of the three macromolecules. The signals at 2135 $cm^{-1}$ ($I_{2135}$), 2185 $cm^{-1}$ ($I_{2185}$), and 2210 $cm^{-1}$ ($I_{2210}$) are linear combination of lipid, protein, and DNA concentrations ($CD_L$, $CD_P$, and $CD_{DNA}$) with coefficients $a_L$, $a_P$, $a_{DNA}$, $b_L$, $b_P$, $b_{DNA}$, $c_L$, $c_P$, and $c_{DNA}$ as the following:

$$\begin{pmatrix} I_{2135} \\ I_{2185} \\ I_{2210} \end{pmatrix} = \begin{pmatrix} a_L & a_P & a_{DNA} \\ b_L & b_P & b_{DNA} \\ c_L & c_P & c_{DNA} \end{pmatrix} \begin{pmatrix} CD_L \\ CD_P \\ CD_{DNA} \end{pmatrix}$$

Unmixng coefficients were obtained from the spectra of D-labeled cellular extracts from $D_2O$-treated HeLa cells (Supplementary Figure 2A). $a_L$, $b_P$, and $c_{DNA}$ were set to 1, and the rest coefficients were scaled to their relative values. Substitute the coefficients with their values, we obtained:

$$\begin{pmatrix} I_{2135} \\ I_{2185} \\ I_{2210} \end{pmatrix} = \begin{pmatrix} 1 & 0.38 & 0.29 \\ 0.48 & 1 & 1.08 \\ 0.24 & 0.71 & 1 \end{pmatrix} \begin{pmatrix} CD_L \\ CD_P \\ CD_{DNA} \end{pmatrix},$$

from which we derived:

$$\begin{pmatrix} CD_L \\ CD_P \\ CD_{DNA} \end{pmatrix} = \begin{pmatrix} 1.31 & -0.98 & 0.67 \\ -1.24 & 5.21 & -5.27 \\ 0.56 & -3.47 & 4.58 \end{pmatrix} \begin{pmatrix} I_{2135} \\ I_{2185} \\ I_{2210} \end{pmatrix}$$

where $CD_L$, $CD_P$, and $CD_{DNA}$ are unmixed D-labeled lipid, protein, and DNA signal intensity, respectively.

**$CD_L/CD_P$ unmixing**

Since unmixed $CD_{DNA}$ signal is very weak in non-dividing cells and in most of the tissues we imaged, we developed a simplified unmixing method that focused on separating $CD_L$ and $CD_P$ signals. The unmixing of $CH_L$ lipids and $CH_P$ protein signals was performed as previously described[8].

We simplified the three-component unmixing equation and removed the $CD_{DNA}$ variable to obtain the following equation:

$$I_{2135} = a_L \cdot CD_L + a_P \cdot CD_P$$

$$I_{2185} = b_L \cdot CD_L + b_P \cdot CD_P$$

from which we can derive:

$$CD_L = \frac{a_P \cdot I_{2185} - b_P \cdot I_{2135}}{a_P \cdot b_L - a_L \cdot b_P}$$

$$CD_P = \frac{b_L \cdot I_{2135} - a_L \cdot I_{2185}}{a_P \cdot b_L - a_L \cdot b_P}$$

For $CD_L/CD_P$ unmixing, we used the spectra of *in situ* CD lipid and protein signals to measure the coefficients. From the spontaneous Raman spectra of pure D-labeled protein signal (signals after 24-hour methanol wash) and pure D-labeled lipid signal (signals removed by methanol wash—subtracting methanol-resistant signal from total signal). All signals were first normalized to the phenylalanine peak at 1004 $cm^{-1}$, which stayed constant during methanol wash, and then normalized to the peaks of pure protein and lipid signals, respectively (Supplementary Figure 2). Thus, $a_L$ and $b_P$ were set to 1; the relative intensity of protein bleed-through signal in the lipid channel ($a_P$) and the relative intensity of lipid bleed-through signal in the protein channel ($b_L$) were measured on the spectra. For example, in xenograft tumor tissues (Supplementary Figure 2C),

$$a_L = 1;\ a_P = 0.40;\ b_L = 0.51;\ b_P = 1;$$

Thus,

$$CD_L = 1.25 \cdot I_{2135} - 0.50 \cdot I_{2185}$$

$$CD_P = 1.25 \cdot I_{2185} - 0.64 \cdot I_{2135}$$

Using the same method, we also obtained pure D-labeled protein and lipid signals from the mouse pancreas and brain tissues and calculated their specific unmixing coefficients. Data from the three types of tissues were combined to generate a set of average coefficients as the following:

$$a_L = 1;\ a_P = 0.48 \pm 0.17;\ b_L = 0.42 \pm 0.07;\ b_P = 1.$$

Among the multiple tissues we analyzed, the CD lipid spectrum showed a consistent shape, but CD protein had varying levels of bleed-through into CDL channel across different tissue types (Supplementary Figure 3C), which possibly resulted from tissue-specific incorporation of deuterium into various NEAAs and/or different NEAA composition of D-labeled proteins. One extreme example is the pyramidal neurons: their cell body showed almost no SRS signal at 2135 $cm^{-1}$ but strong signal at 2185 $cm^{-1}$, suggesting almost no protein bleed-through into CDL channel. Thus, when applying the unmixing algorithm to cells and tissues previously uncharacterized, we tested $a_P$ from 0.31~0.65 until the nucleus was deprived of $CD_L$ signal after unmixing. In this study, the following equations were used for $CD_L/CD_P$ signal unmixing of different tissues.

| | $CD_L$ unmixing | $CD_P$ unmixing |
|---|---|---|
| Cos-7 and HeLa cells | $CD_L = 1.25 \cdot I_{2135} - 0.39 \cdot I_{2185}$ | $CD_P = 1.25 \cdot I_{2185} - 0.52 \cdot I_{2135}$ |
| Glioblastoma and Colon tumor xenograft | $CD_L = 1.25 \cdot I_{2135} - 0.50 \cdot I_{2185}$ | $CD_P = 1.25 \cdot I_{2185} - 0.64 \cdot I_{2135}$ |
| Zebrafish tissues | $CD_L = 1.25 \cdot I_{2135} - 0.50 \cdot I_{2185}$ | $CD_P = 1.25 \cdot I_{2185} - 0.52 \cdot I_{2135}$ |
| *C. elegans* | $CD_L = 1.25 \cdot I_{2135} - 0.70 \cdot I_{2185}$ | $CD_P = 1.25 \cdot I_{2185} - 0.52 \cdot I_{2135}$ |
| Mouse sebaceous glands and adipose tissues | $CD_L = 1.25 \cdot I_{2135} - 0.39 \cdot I_{2185}$ | $CD_P = 1.25 \cdot I_{2185} - 0.81 \cdot I_{2135}$ |
| Mouse internal capsule and cortex | $CD_L = 1.25 \cdot I_{2135} - 0.39 \cdot I_{2185}$ | $CD_P = 1.25 \cdot I_{2185} - 0.70 \cdot I_{2135}$ |
| Mouse liver, pancreas, and muscle tissues | $CD_L = 1.25 \cdot I_{2135} - 0.35 \cdot I_{2185}$ | $CD_P = 1.25 \cdot I_{2185} - 0.60 \cdot I_{2135}$ |

#### Image processing

We used Olympus FluoView 4.1a scanning software to acquires images and ImageJ to assign color, calculate ratiometric values, and overlay images. $CD_L/CH_L$ and $CD_P/CH_P$ ratios were used to show the proportion of newly synthesized lipids and proteins to total macromolecules, as an indication of relative metabolic rate.

For example,

$$\frac{CD_L}{CH_L} = \frac{a * [C - D]}{b * [C - H]} = \frac{a * [\text{new lipid}] * G}{b * ( [\text{new lipid}] * (1 - G) + [\text{old lipid}] )}$$

, where $[C – D]$ and $[C – H]$ are the concentration of D-labeled and H-labeled lipids, respectively; $a$ and $b$ are converting factors from lipid concentrations to SRS signals intensity; [old lipid] is the amount of lipids that pre-existed before $D_2O$ probing and remained at the time of imaging (not being degraded); [new lipid] is the amount of newly synthesized lipids and $G$ is the proportion of D-labeled lipids to the newly synthesized lipids.

If we define [total lipid] as the amount of total lipid after the probing period and at the time of imaging, [total lipid] = [new lipid] + [old lipid]. Then the above equation can be written as:

$$\frac{CD_L}{CH_L} = \frac{a}{b} \cdot \frac{G}{\frac{[\text{total lipid}]}{[\text{new lipid}]} - G}$$

Since the upper limit of $G$ is ~0.2 ($D_2O$ enrichment) in all our experiment, much smaller than [total lipid]/[new lipid], the $CD_L/CH_L$ can be approximated as:

$$\frac{CD_L}{CH_L} = \frac{a}{b} \cdot G \cdot \frac{[\text{new lipid}]}{[\text{total lipid}]}$$

Thus, $CD_L/CH_L$ is linear to the proportion of the amount of newly synthesized lipids to total lipids and therefore a ratiometric measurement of lipid synthesis rate.

#### References


1. Shen, Y., Xu, F., Wei, L., Hu, F. & Min, W. Live-cell quantitative imaging of proteome degradation by stimulated Raman scattering. *Angew Chem Int Ed Engl* **53**, 5596-5599 (2014).
2. Wei, L. et al. Live-cell imaging of alkyne-tagged small biomolecules by stimulated Raman scattering. *Nat Methods* **11**, 410-412 (2014).
3. Brenner, S. The genetics of Caenorhabditis elegans. *Genetics* **77**, 71-94 (1974).
4. Choi, J. et al. FoxH1 negatively modulates flk1 gene expression and vascular formation in zebrafish. *Dev Biol* **304**, 735-744 (2007).
5. Westerfield, M. The zebrafish book. A guide for the laboratory use of zebrafish (Danio rerio). 4th ed. (Univ. of Oregon Press, Eugene; 2000).
6. Rodriguez-Contreras, A., Shi, L. & Fu, B.M. A method to make a craniotomy on the ventral skull of neonate rodents. *J Vis Exp* (2014).
7. Lu, F.K. et al. Label-free DNA imaging in vivo with stimulated Raman scattering microscopy. *Proc Natl Acad Sci U S A* **112**, 11624-11629 (2015).

8. Lu, F.K. et al. Multicolor stimulated Raman scattering (SRS) microscopy. *Mol Phys* **110**, 1927-1932 (2012).
