## Supplementary Materials for "Optical Imaging of Metabolic Dynamics in Animals"

**Supplementary Figures and Movies**

**Supplementary Figure 1. Toxicity of $D_2O$ on cells and *C. elegans*.**

**Supplementary Figure 2. The development of $CD_L/CD_P$ unmixing algorithm.**

**Supplementary Figure 3. Imaging lipogenesis in animals.**

**Supplementary Figure 4. Imaging *de novo* protein biosynthesis.**

**Supplementary Figure 5. DO-SRS tracks the metabolism of specific lineage during embryonic development in zebrafish.**

**Supplementary Movie 1. *In vivo* SRS imaging of D-labeled lipids in the sebaceous glands of a living mouse at 2135 $cm^{-1}$**

**Supplementary Movie 2. *In vivo* SRS imaging of pre-existing lipids in the sebaceous glands of a living mouse at 2845 $cm^{-1}$**

**Supplementary Movie 3. SRS live imaging of D-labeled lipids of moving *C. elegans* larvae at 2135 $cm^{-1}$**

**Supplementary Movie 4. SRS live imaging of pre-existing lipids of moving *C. elegans* larvae at 2845 $cm^{-1}$**

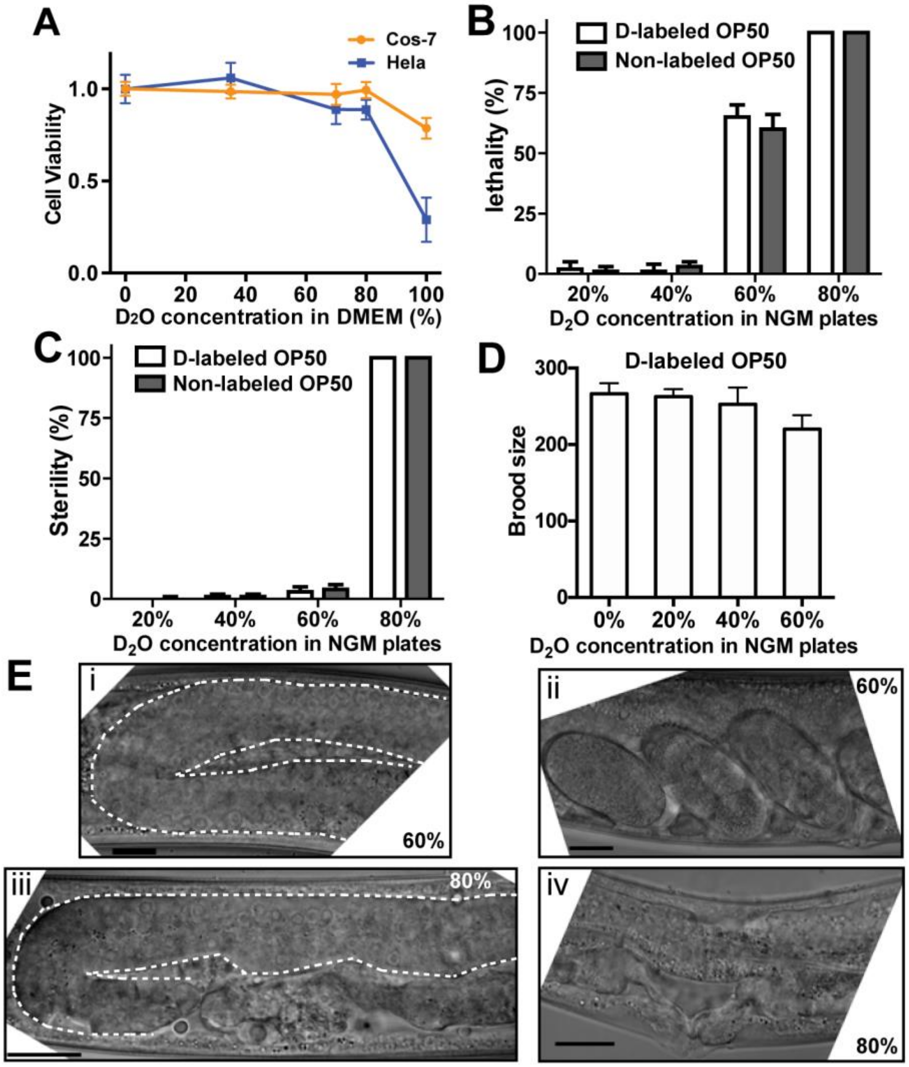


**Supplementary Figure 1. Toxicity of $D_2O$ on cells and *C. elegans***. (A) CellTiter-Glo Luminescent Cell Viability Assay (Promega), which quantifies ATP production, was performed on Cos-7 and HeLa cells grown in DMEM made of different concentration of $D_2O$ for 48 hours. (B) Lethality was determined by placing hypochlorite-prepared eggs onto NGM plates made of different concentration of $D_2O$ and calculating the percentage of eggs that hatched into normal, moving larva. D-labeled OP50 refers to overnight grown culture of *E. coli* OP50 that were seeded onto the $D_2O$-containing NGM plates 24 hours before the experiment; non-labeled OP50 refers to the experimental setup, in which bacteria were killed by UV immediately after being placed onto the $D_2O$ plates and thus was not labeled by deuterium. (C) Sterility was determined as the percentage of fourth stage larva (L4) that became sterile after being transferred to $D_2O$ plates. (D) Brood size, which reflects the effects on meiosis, was determined by counting the total number of viable progeny one animal produced. (E) At 60% $D_2O$ concentration, the gonad developed normally (i) and normal embryos were formed (ii) in animals that developed from L4 to adults on $D_2O$ plates. With 80% $D_2O$, gonad development was disrupted and proliferation of the germ cells appeared to be affected (iii) and no embryos were formed in the uterus (iv). Scale bar = 20 μm.

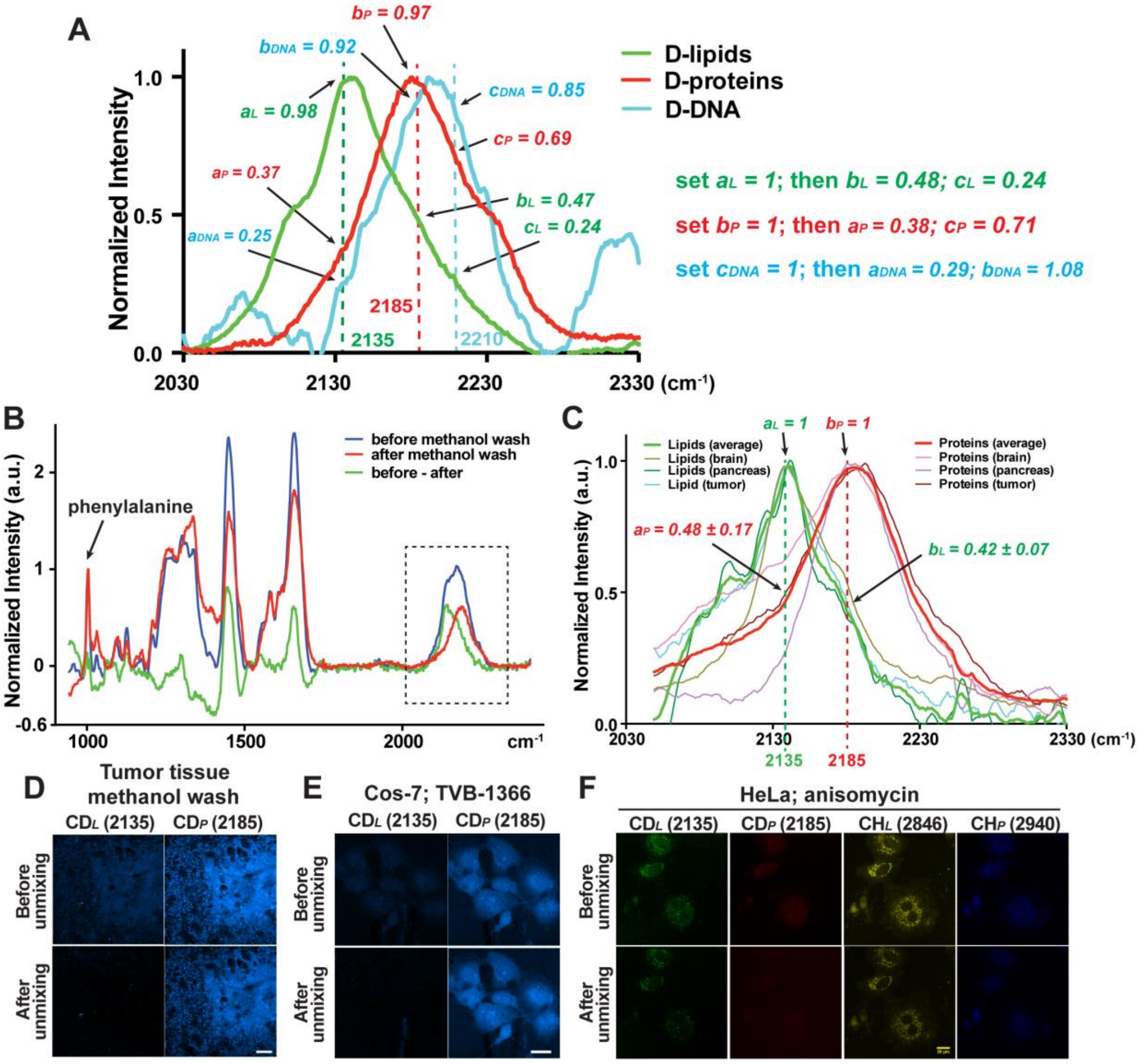


**Supplementary Figure 2. The development of three-component unmixing algorithm.** (A) Spontaneous Raman signal of extracted lipids, proteins, and DNA from $D_2O$-treated HeLa cells. Signals for each type of macromolecules were normalized to their own peaks. Unmixing coefficients were first measured from the spectra and then adjusted by setting $a_L$, $b_P$, and $c_{DNA}$ to 1 and scaling the other parameters accordingly. (B) Colon tumor tissues (from mice that drank 25% $D_2O$ for 15 days) before and after methanol wash. Pure deuterium-labeled protein signals (red) were assumed as the methanol-resistant signal, and pure deuterium-labeled lipid signals (green) were calculated as the signal difference before and after methanol wash. All signals were normalized to the phenylalanine peak. (C) Similar methanol wash and signal normalization methods were applied to mouse brain and pancreas tissues, in addition to tumor tissues. The average, normalized pure protein and lipid signal intensities from the three tissues were calculated; mean ± SD for the coefficients were shown. (D) SRS images of Cos-7 cells that were grown in 75% $D_2O$ DMEM for 24 hours before and after methanol wash and before and after unmixing calculation. (E) SRS images of methanol-washed colon tumor tissues (taken from tumor-bearing mice that drank 25% $D_2O$ for 15 days) before and after unmixing ($a_L$ = 1; $a_P$ = 0.40; $b_P$ = 1; $b_L$ =

0.51). Unmixing was also applied to Cos-7 cells grown in 70% $D_2O$ DMEM containing 10 nM TVB-3166 for 24 hours (F) and HeLa cells grown in 70% $D_2O$ DMEM containing 1 μM anisomycin for 24 hours (G). Unmixing equations can be found in the Materials and Methods. Related to Figure 3.

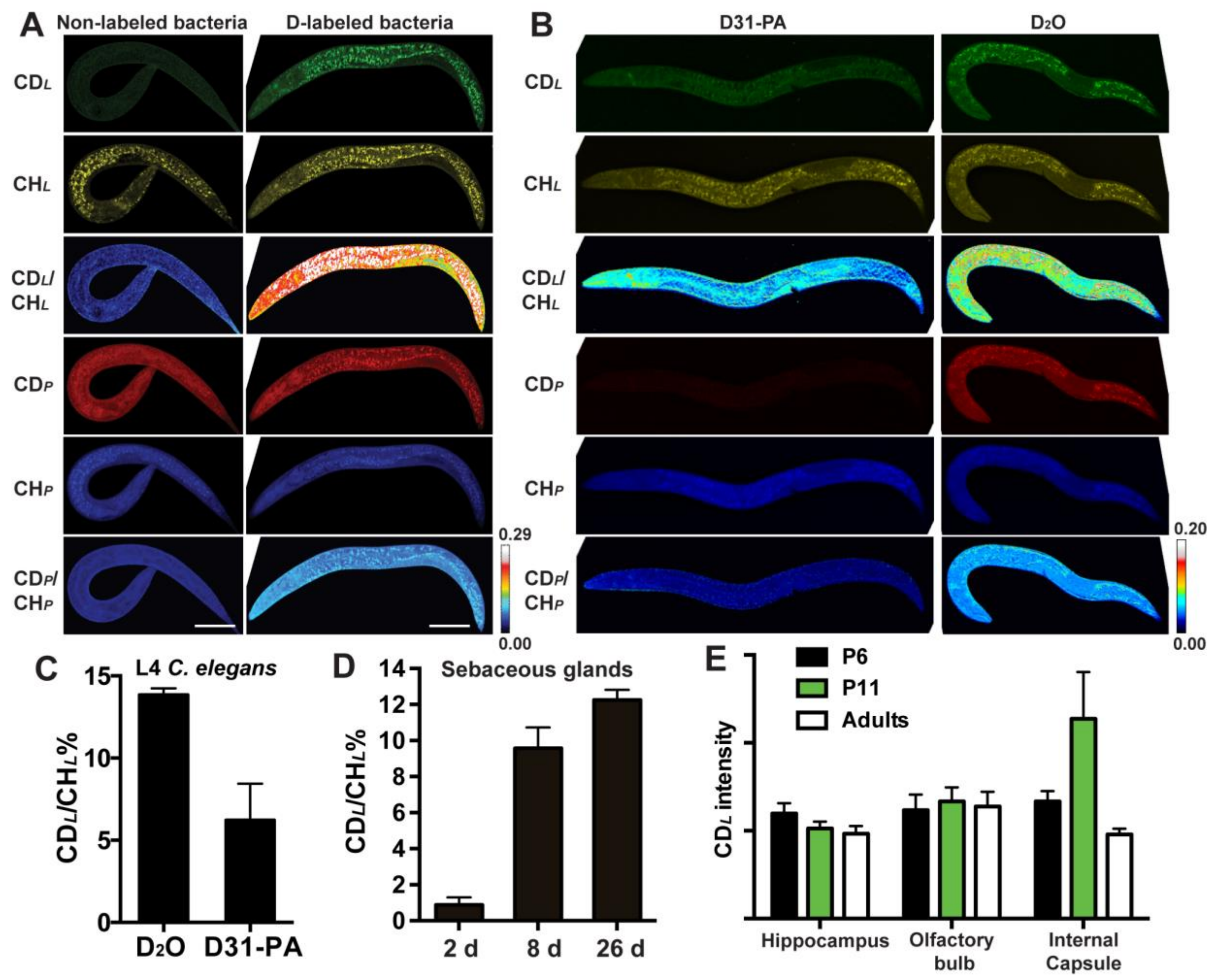


**Supplementary Figure 3. Imaging lipogenesis in animals.** (A) Hypochlorite-prepared eggs of *C. elegans* were placed onto 20% $D_2O$ NGM plates pre-seeded with *E. coli* OP50 and third stage larva grown from those eggs were imaged. For the live bacteria group, OP50 grew on the $D_2O$ plates for 24 hours at room temperature before eggs were placed; for the dead bacteria group, bacteria cells were killed by UV immediately after being seeded onto the $D_2O$ plates, and 24 hours later, eggs were placed onto the plate. Quantification is shown in Figure 4E. (B) OP50 bacterial culture was mixed with 4 mM *D31*-palmitic acid and then seeded onto NGM plates that contains 100% $H_2O$. *C. elegans* eggs were placed onto those plates and as controls 20% $D_2O$ plates with OP50 seeded the day before. 48 hours later, L4 animals were imaged. (C) Comparison of the $CD_L/CH_L$ ratio for animals fed with D31-PA and the ones grown on $D_2O$ plates. (D) Quantifications of lipid synthesis rate in the sebaceous glands. The ratio of $CD_L$ mean intensity to $CH_L$ mean intensity across an entire gland unit were quantified for tissues taken from mice that drank 25% $D_2O$ for 2 or 8 or 26 days. (C) Quantification of myelination activities in different brain regions. Several axon fiber-rich region of the brain were imaged for P6 and P11 pups and adults. Pups were fed on milk for 6 days from the mothers that drank 25% $D_2O$; adults drank 25% $D_2O$ for 9 days. $CD_L$ signal were quantified as the mean intensity of the entire image. Related to Figure 4.

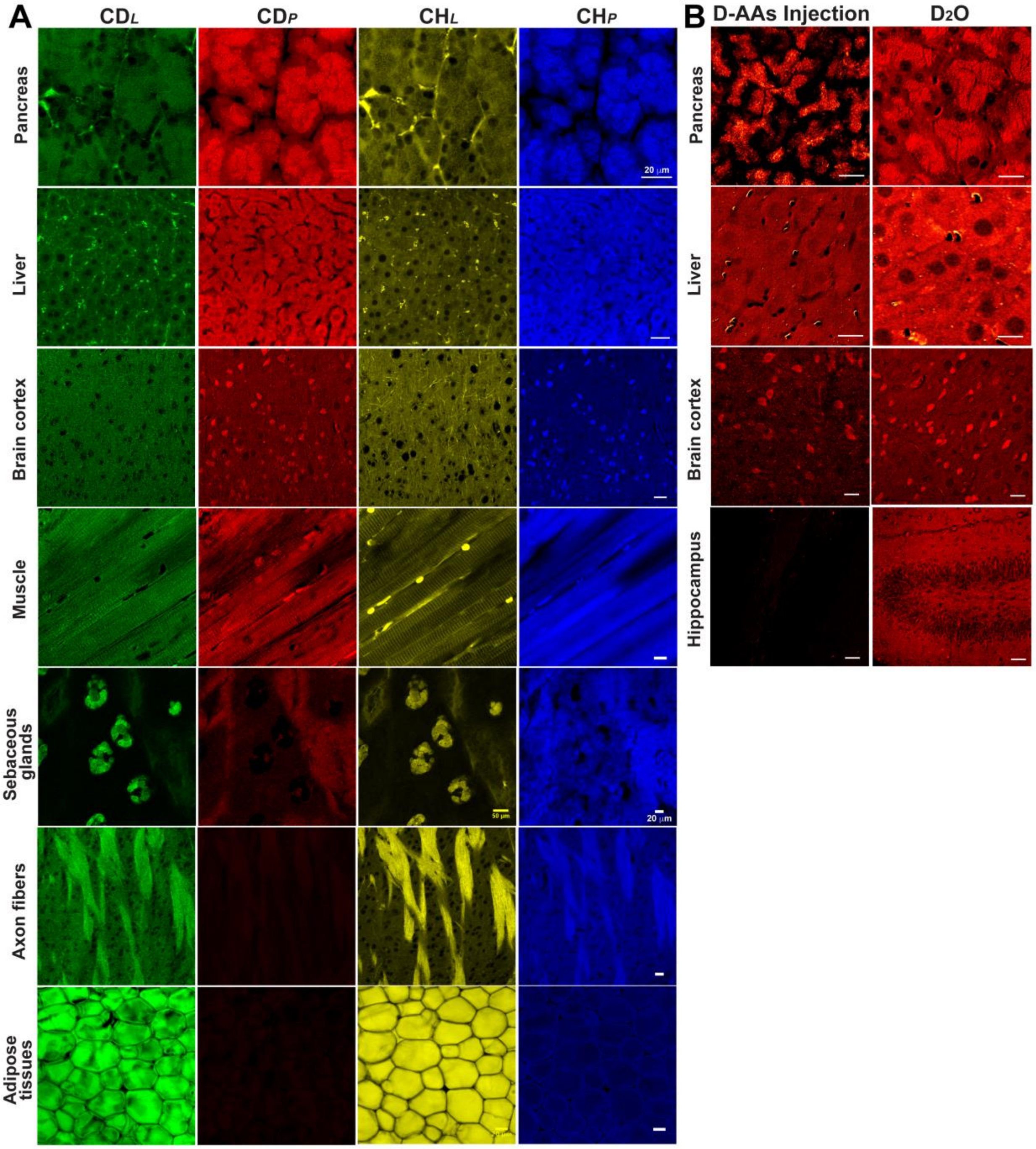


**Supplementary Figure 4. Imaging *de novo* protein biosynthesis.** (A) Various protein-rich tissues, including pancreas, liver, brain cortex, and muscle, were collected from adult mice that drank 25% $D_2O$ for 20 days and then imaged. Lipid-rich sebaceous glands and adipose tissues were harvested from adult mice that drank 25% $D_2O$ for 8 and 15 days, respectively. Myelinated axon fibers of internal capsule were harvested from P11 mouse pups that were fed on milk produced by mother mice drinking 25% $D_2O$ for 6 days before imaging. (B) $CD_P$ signals of various organs from mice that were injected with D-labeled amino acids (d-AAs) *via* carotid artery and mice that drank 25% $D_2O$ for 8 days. Scale bars = 20 μm. Related to Figure 5.

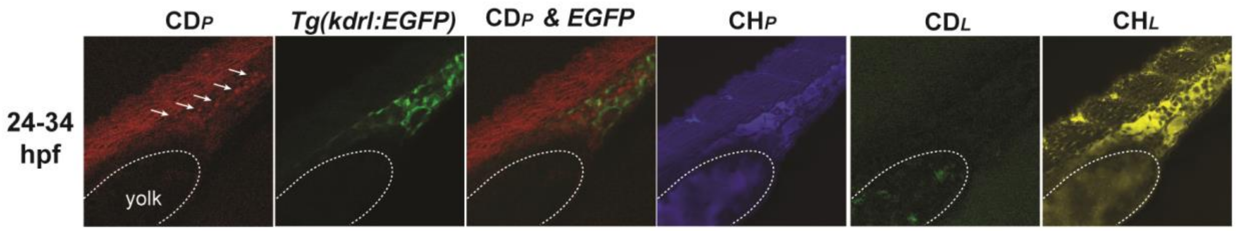


**Supplementary Figure 5. DO-SRS tracks the metabolism of specific lineage during embryonic development in zebrafish.** SRS signal and the colocalization with fluorescence from *Tg(kdrl:EGFP)* reporter in zebrafish embryos that were incubated in egg solution containing 20% $D_2O$ from 24 to 34 hpf. Dashed curves outline the yolk sac extension (Y). A few cells with strong $CD_P$ signal (arrows) were also labeled by GFP and are likely differentiating angioblasts. Related to Figure 6.
